# Targeting FSP1 triggers ferroptosis in lung cancer

**DOI:** 10.1101/2025.08.07.668766

**Authors:** Katherine Wu, Alec J Vaughan, Jozef P Bossowski, Yuan Hao, Aikaterini Ziogou, Seon Min Kim, Tae Ha Kim, Mari N Nakamura, Ray Pillai, Mariana Mancini, Sahith Rajalingam, Mingqi Han, Toshitaka Nakamura, Lidong Wang, Suckwoo Chung, Diane Simeone, David Shackelford, Yun Pyo Kang, Marcus Conrad, Thales Papagiannakopoulos

**Affiliations:** Department of Pathology, New York University Grossman School of Medicine, New York, NY, USA; Vilcek Institute of Graduate Biomedical Sciences, New York University Grossman School of Medicine, New York, NY, USA; Laura and Isaac Perlmutter Cancer Center, New York University Langone Health, New York, NY, USA; College of Pharmacy and Research Institute of Pharmaceutical Sciences, Seoul National University, Seoul, Korea; Division of Pulmonary and Critical Care Medicine, Department of Medicine, VA New York Harbor Healthcare System, New York, USA; Division of Pulmonary, Critical Care, and Sleep Medicine, Department of Medicine, New York University Grossman School of Medicine, New York, USA; Pulmonary and Critical Care Medicine, David Geffen School of Medicine (DGSOM), University of California Los Angeles (UCLA), Los Angeles, CA, USA; Jonsson Comprehensive Cancer Center, UCLA, Los Angeles, CA, USA; Moores Cancer Center, University of California, San Diego, La Jolla, CA, USA; Institute of Metabolism and Cell Death, Molecular Targets and Therapeutics Center, Helmholtz Munich, Neuherberg, Germany; Translational Redox Biology, Technical University of Munich (TUM), TUM Natural School of Sciences, Garching, Germany

**Keywords:** lung cancer, ferroptosis, FSP1, GPX4

## Abstract

Pre-clinical and clinical studies have demonstrated how dietary antioxidants or mutations activating antioxidant metabolism promote cancer, highlighting a central role oxidative stress in tumorigenesis. However, it is unclear if oxidative stress ultimately increases to a point of cell death. Emerging evidence indicates that cancer cells are susceptible to ferroptosis, a form of cell death triggered by uncontrolled lipid peroxidation^1–3^. Despite broad enthusiasm about harnessing ferroptosis as a novel anti-cancer strategy, whether ferroptosis is a barrier to tumorigenesis and if it can be leveraged therapeutically remains unknown^4,5^. Using genetically-engineered mouse models (GEMMs) of lung adenocarcinoma (LUAD), we performed tumor specific loss-of-function studies of the two key ferroptosis suppressors, *glutathione peroxidase 4* (*Gpx4*)^6,7^ and *ferroptosis suppressor protein 1* (*Fsp1*)^8,9^, and observed increased lipid peroxidation and robust suppression of tumorigenesis, suggesting that lung tumors are highly sensitive to ferroptosis. Furthermore, across multiple pre-clinical models, we found that FSP1 was required for ferroptosis protection *in vivo*, but not *in vitro*, underscoring a heightened need to buffer lipid peroxidation under physiological conditions. Lipidomic analyses revealed that Fsp1-knockout (Fsp1^KO^) tumors had an accumulation of lipid peroxides, and inhibition of ferroptosis with genetic, dietary, or pharmacological approaches effectively restored the growth of Fsp1^KO^ tumors *in vivo*. Unlike *GPX4*, *FSP1* expression was prognostic for disease progression and poorer survival in LUAD patients, highlighting its potential as a viable therapeutic target. To this end, we demonstrated that pharmacologic inhibition of FSP1 had significant therapeutic benefit in pre-clinical lung cancer models. Our studies highlight the importance of ferroptosis suppression *in vivo* and pave the way for FSP1 inhibition as a therapeutic strategy in lung cancer patients.

## MAIN TEXT

Ferroptosis is a non-apoptotic, oxidative stress-dependent mechanism of cell death that is uniquely distinguished by lipid peroxidation of polyunsaturated fatty acids of membrane phospholipids (PUFA-PL)^1,10^. Lipid peroxidation involves the formation of lipid and lipid peroxyl radicals which, in auto-oxidative propagation reactions, generate lipid hydroperoxides. Aberrant, unrestricted lipid peroxidation results in altered membrane integrity, cell swelling, and ultimately membrane rupture^11,12^. In recent years, growing interest in ferroptosis has elucidated that cells depend on several enzymes to protect against ferroptosis^6–9,13–15^, of which Glutathione Peroxidase 4 (GPX4) and Ferroptosis Suppressor Protein 1 (FSP1) are the most notable. GPX4 catalyzes the reduction of PUFA-PL hydroperoxides to non-toxic alcohols. FSP1 catalyzes the reduction of extramitochondrial Coenzyme Q_10_ (CoQ), a highly potent lipid radical-trapping antioxidant (RTA). On the other hand, cells can be sensitized to ferroptosis through PUFA-PL composition^16^ and biosynthesis by Long-chain Acyl-CoA Synthase 4 (ACSL4)^17–19^. Apart from these cellular pathways, exogenous lipid RTAs, such as liproxstatin-1 (LIP1), ferrostatin-1 (FER1), Vitamin E (VitE), and Vitamin K (VitK), have also been shown to specifically inhibit ferroptosis^1,6,20–22^.

Emerging evidence indicates that cancer cells, including drug-resistant cells that have adopted a mesenchymal state^2,3^, are highly sensitive to lipid peroxidation *in vitro*, garnering excitement about harnessing ferroptosis as a novel anti-cancer strategy. Though there have been extensive studies characterizing GPX4 and more recently FSP1 in various diseases^4,23^, whether ferroptosis constitutes a barrier to tumorigenesis and the functional roles of GPX4 and FSP1 in *in vivo* cancer models remain poorly characterized. To date, FSP1 has been shown to be a critical ferroptosis suppressor only in the absence of GPX4^8,9,24^. However, given the high toxicity, poor selectivity, and low to limited bioavailability of GPX4 inhibitors *in vivo*^25,26^ as well as the only recent development of FSP1 inhibitors, the majority of which do not have *in vivo* efficacy^9,24,27–30^, much work remains to demonstrate that ferroptosis induction could indeed be a therapeutic strategy for cancer^5^.

Here we show that genetic knockout of either Gpx4 or Fsp1 in genetically-engineered mouse models (GEMMs) of autochthonous Kras-driven lung adenocarcinoma (LUAD)^31–35^ results in dramatic restriction of lung tumorigenesis. We demonstrate that disruption of Fsp1, often considered a backup axis of ferroptosis suppression, is unexpectedly sufficient to trigger exhibit enhanced lipid peroxidation in lung tumors. We confirm that Fsp1 deletion across several human cancer cell lines with various oncogenic drivers, co-mutations, and tissue lineages consistently lead to *in vivo* tumor restriction. As ferroptosis can be functionally defined as a modality of cell death resulting from elevated lipid peroxides and rescued only with inhibitors of lipid peroxidation, we demonstrate that Fsp1-knockout (Fsp1^KO^) tumors have increased oxidized PUFA-PLs and that administration of lipid RTAs, as well as genetic loss of Acsl4, effectively restores lung tumorigenesis. Finally, we report that FSP1 is upregulated as LUAD tumors progress and show that FSP1 inhibition suppresses tumor growth and prolongs survival in multiple pre-clinical cancer models. Thus, our work utilizes novel *in vivo* models to characterize and test ferroptosis induction, particularly via FSP1 perturbation, as a therapeutic approach for lung cancer.

### GPX4 prevents ferroptosis in lung tumors

GPX4 is considered the primary mediator of ferroptosis suppression that has been shown to be essential across a plethora of murine and human cancer cells^7,26,36,37^. Therefore, we sought to interrogate whether GPX4, and more broadly ferroptosis suppression, is a requirement for lung tumorigenesis *in vivo*. Using a well-established GEMM of LUAD (*Kras^LSL-G12D/+^, Tp53^fl/fl^*, *Rosa26*^LSL-Cas9/LSL-Cas9^), we initiated Kras^G12D/+^, Tp53^−/−^ (KP) tumors by intratracheal delivery of lentiviruses expressing CRE-recombinase and dual sgRNAs^38^ against either Gpx4 (sg*Gpx4*) or non-targeting control (sg*Neo*), enabling CRISPR/Cas9-mediated Gpx4-knockout (Gpx4^KO^) or wildtype (WT) control tumors, respectively. Concurrently, we treated a cohort of animals with each tumor genotype with Liproxstatin-1 (LIP1), a potent lipid RTA and ferroptosis inhibitor, throughout the entire course of tumorigenic progression (Fig. 1a). Tumor-specific Gpx4 deletion (sg*Gpx4*) led to a significant decrease in lung tumor burden that was not observed in sg*Gpx4* tumor-bearing mice treated with LIP1 (Fig. 1b-d). Efficient Gpx4 deletion was verified by immunohistochemistry (IHC), with less than 15% of tumors in the in sg*Gpx4* group still staining positive for Gpx4 (Extended Data Fig. 1a). IHC for 4-hydroxy-2-noneal (4-HNE), a marker for lipid peroxidation in tissues (Extended Data Fig. 1b)^20^, revealed significantly higher levels of lipid peroxidation in sg*Gpx4* tumors compared to control sg*Neo* lung tumors, which was blunted with LIP1 treatment (Fig. 1e, f).

**Fig. 1:**
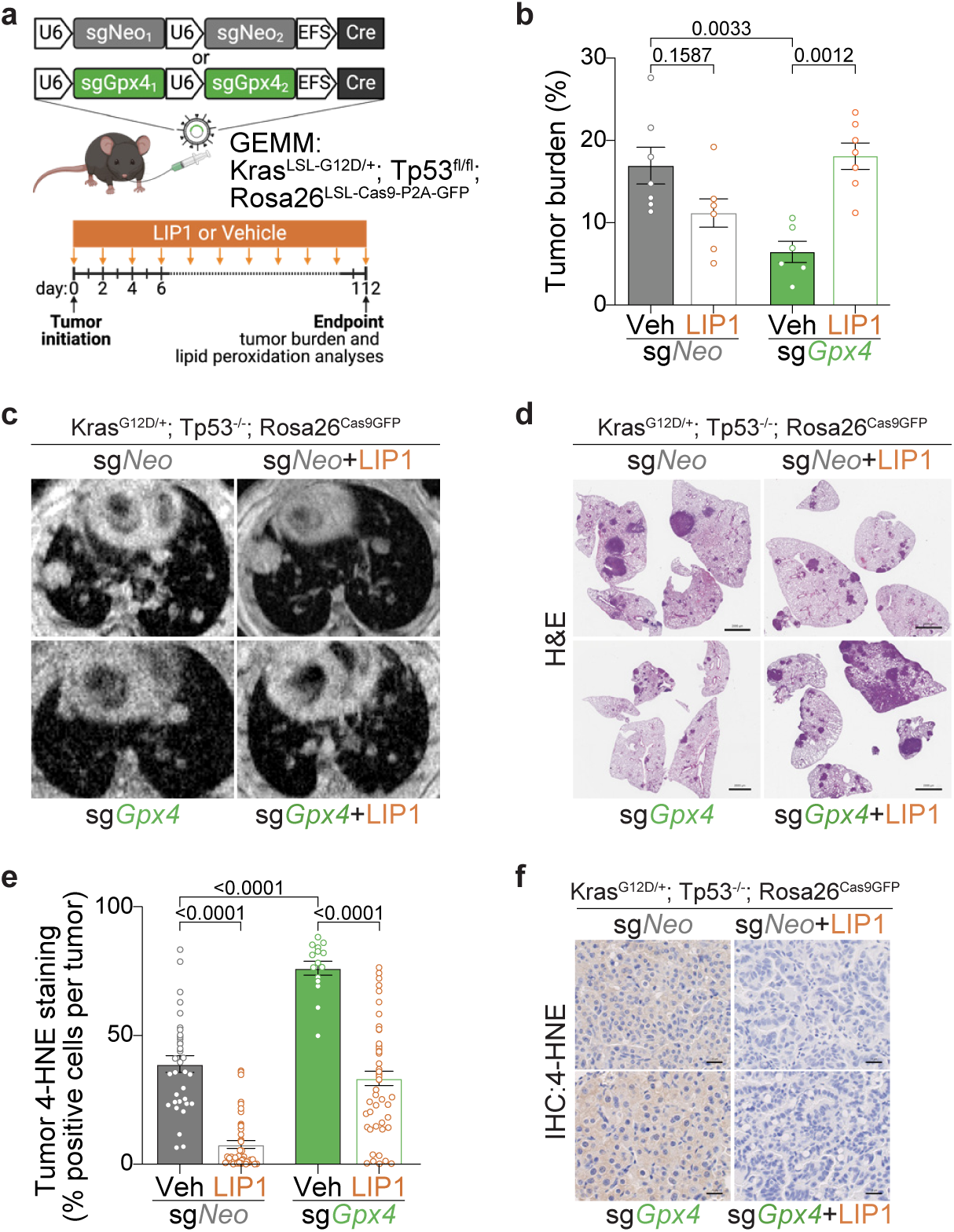
Gpx4 loss triggers ferroptosis in lung tumors. **(a)** Schematic of KP LUAD GEMMs intratracheally infected with pUSEC lentiviruses containing dual sgRNAs targeting Neo (control; n=13) or Gpx4 (n=13). Mice were dosed with Liproxstatin-1 (LIP1; sg*Neo* n=6, sg*Gpx4* n=7) or Vehicle (Veh; sg*Neo* n=6, sg*Gpx4* n=7) every other day starting from tumor initiation to experiment endpoint. **(b)** Tumor burden of KP LUAD tumors with control (sg*Neo*) or Gpx4 (sg*Gpx4*) knockout, treated with either Veh or LIP1. **(c)** Representative MRI of KP LUAD tumors with control (sg*Neo*) or Gpx4 (sg*Gpx4*) knockout, treated with either Veh or LIP1. **(d)** Representative H&E of KP LUAD tumors with control (sg*Neo*) or Gpx4 (sg*Gpx4*) knockout, treated with either Veh or LIP1. Scale bars: 2000µm. **(e)** Tumor 4-HNE expression by IHC of KP LUAD tumors with control (sg*Neo*) or Gpx4 (sg*Gpx4*) knockout, treated with either Veh or LIP1. **(f)** Representative 4-HNE IHC staining from (e). Scale bars: 20µm. Data are represented as mean values, error bars represent SEM, significance determined via one-way ANOVA with multiple comparisons (panels b, e).

Similar to our *in vivo* observations, CRISPR/Cas9-mediated Gpx4^KO^ in KP LUAD cells resulted in a near complete loss of viability and clonogenicity that was fully rescued by the addition of LIP1 (Extended Data Fig. 1c). Targeted lipidomics to profile oxidized phosphatidylcholine (PCs) and phosphatidylethanolamine (PEs) species in KP LUAD cells treated with RSL3, a covalent Gpx4 inhibitor and ferroptosis inducer, revealed a higher abundance of oxidized phospholipids in RSL3-treated cells that again was decreased in cells treated with concomitant LIP1 (Extended Data Fig. 1d). On the other hand, Gpx4 overexpression (Gpx4^OE^) in KP LUAD cells had no impact on cell viability or clonogenicity basally (Extended Data Fig. 1e) but did confer greater resistance to ferroptosis *in vitro* (Extended Data Fig. 1f). Similarly, supplementation of KP LUAD cells with sodium selenite (Na_2_SeO_3_) to increase translation of Gpx4^39^, a selenoprotein, resulted in a corresponding boost in Gpx4 expression and enhanced ferroptosis resistance *in vitro* (Extended Data Fig. 1g). Taken together, these results demonstrate that Gpx4 is essential for lung tumors to evade ferroptosis and provide evidence that ferroptosis induction may be an effective anti-cancer strategy.

### Unlike GPX4, FSP1 is upregulated in LUAD

Given our observation that Gpx4 was required for lung tumorigenesis in a GEMM that closely recapitulates human LUAD development and progression, we next asked whether there was clinical evidence suggesting that ferroptosis regulators in general are altered in lung cancer^40^. Although GPX4 expression was modestly increased in *KRAS-*mutant LUAD tumors compared to normal lung tissue, there was no correlation with patient tumor stage or overall patient survival (Extended Data Fig. 1h, i). Concordantly, Gpx4 protein levels also did not change during autochthonous LUAD progression in KP GEMM tumors (Extended Data Fig. 1j).

Since GPX4 expression did not appear to correlate with any clinical prognostic factors in *KRAS-*mutant LUAD patients, we next investigated FSP1, a second axis of ferroptosis surveillance (Fig. 2a). We found that FSP1 expression was robustly increased in *KRAS*-mutant LUAD patient tumors compared to normal lung and exhibited a positive correlative trend with higher tumor stages (Extended Data Fig. 2a). Additionally, high FSP1 expression correlated with poor survival in *KRAS-*mutant LUAD patients (Fig. 2b). Unlike Gpx4, Fsp1 protein levels increased temporally as lung tumors progressed from adenomas to adenocarcinomas in KP GEMMs (Figs. 2c, d). Altogether, these findings suggest that FSP1 may constitute a major suppressor of ferroptosis in LUAD and is potentially a more viable therapeutic target than GPX4 for lung cancer patients.

**Fig. 2:**
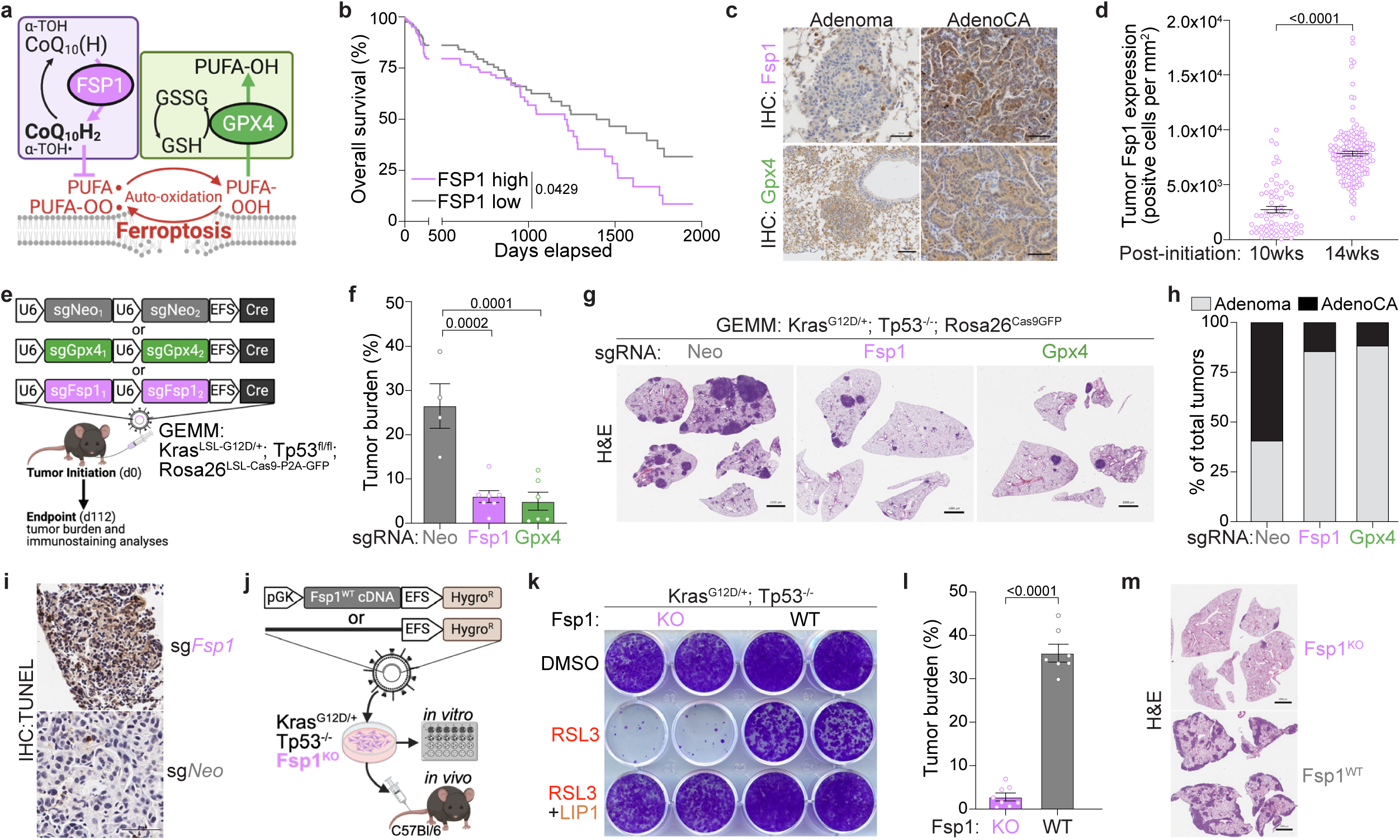
Fsp1 knockout robustly restricts lung tumorigenesis. **(a)** Schematic depicting mechanisms of Fsp1- and Gpx4-mediated buffering against cellular lipid peroxidation and ferroptosis. **(b)** Overall survival of *KRAS*-mutant LUAD patients (n=280) from TCGA, stratified by high versus low primary tumor *FSP1* (*AIFM2*) expression. Median survival for FSP1 high versus low was 1215 days and 1498 days, respectively (hazard ratio 1.51 [1.01-2.25]). **(c)** Representative IHC for Fsp1 and Gpx4 in KP LUAD GEMM adenomas versus adenocarcinomas. Scale bars: 100µm. **(d)** Tumor Fsp1 expression by IHC of KP LUAD GEMM tumors at 10-versus 14-weeks post tumor initiation. **(e)** Schematic of KP LUAD GEMMs intratracheally infected with pUSEC lentiviruses containing double sgRNAs targeting control (Neo; n=4), Fsp1 (n=6), or Gpx4 (n=6). **(f)** Tumor burden of KP LUAD tumors with knockout of either control (Neo), Fsp1, or Gpx4. **(g)** Representative H&E of KP LUAD tumors with knockout of either control (Neo), Fsp1, or Gpx4. Scale bars: 2000µm. **(h)** Relative proportion of KP adenomas versus adenocarcinomas with knockout of either control (Neo), Fsp1, or Gpx4. **(i)** Representative TUNEL staining of KP, Fsp1^KO^ and Fsp1^WT^ orthotopic lung tumors. Scale bars: 50µm **(j)** Schematic depicting generation of isogenic Fsp1-wildtype (Fsp1^WT^) versus Fsp1-knockout (Fsp1^KO^) cells for paralleled *in vitro* assays and *in vivo* syngeneic orthotopic transplantation studies. **(k)** Representative images of crystal violet clonogenic growth assay in isogenic KP, Fsp1^KO^ and Fsp1^WT^ cells treated with RSL3 (0.5µM) ± LIP1 (100nM). **(l)** Tumor burden of KP, Fsp1^KO^ (n=8) and Fsp1^WT^ (n=7) orthotopic lung tumors. **(m)** Representative H&E of KP, Fsp1^KO^ and Fsp1^WT^ orthotopic lung tumors. Scale bars: 2000µm. Data are represented as mean values, error bars represent SEM, significance determined via two-sided student’s t-test (panels d and l), one-way ANOVA with multiple comparisons (panel f) or Kaplan-Meier simple survival analysis (panel b).

Since prior studies have suggested that FSP1 may be regulated by KRAS signaling^41^ to further determine whether FSP1 upregulation was specific to *KRAS-*mutant LUAD, we stratified LUAD patients by major oncogenic mutations. Again, *FSP1* was robustly increased in tumors relative to normal lung, and this upregulation was observed irrespective of driver mutation or *KRAS* variant (Extended Data Fig. 2b, c). Although *FSP1* expression was more robustly upregulated in *KRAS*-mutant tumors compared to *EGFR*-mutant tumors, LUAD tumors with neither mutation also exhibited upregulation of *FSP1* to the same extent as *KRAS*-mutant tumors. Next, we investigated whether MAPK/ERK pathway regulates FSP1 expression. Both *in vitro* pathway inhibition with RMC-042, a pan-RAS inhibitor, in a panel of *KRAS*-mutant LUAD cell lines as well as multi-immunofluorescence (IF) for co-expression of Fsp1 and pErk in KP GEMM tumors revealed a lack of correlation between FSP1 expression and ERK activation (Extended Data Fig. 2d, e). Thus, these data indicate that *FSP1* expression, at least in the context of LUAD, may not be exclusively dependent on KRAS-mediated oncogenic signaling.

Given that patients with *KRAS*-mutant LUAD tumors often have concomitant mutations in *STK11* and *KEAP1*, which portend more advanced disease and poorer prognosis^34,43^, we also wondered whether *FSP1* expression was impacted by these mutations. We observed that *FSP1* was significantly increased in LUAD tumors with co-mutation of either *STK11* or *KEAP1* (Extended Data Fig. 2f). Additionally, treatment of a panel of human LUAD cell lines with an NRF2 activator, KI696, led to significantly increased *FSP1* gene expression (Extended Data Fig. 2g). Of note, these findings are in accordance with recent studies describing a role for NRF2, which is activated by *KEAP1* mutations, in FSP1 upregulation^42^.

### Fsp1 is required for lung tumorigenesis

To systematically investigate the functional role of Fsp1 in lung cancer, we initiated autochthonous KP LUAD tumors, this time with tumor-specific CRISPR/Cas9-mediated deletion of either Fsp1 or Gpx4 (Fig. 2e). Strikingly, we found that genetic deletion of Fsp1 (sg*Fsp1*) in KP LUAD tumors strongly suppressed lung tumorigenesis, resulting in significantly decreased tumor burden to the same extent as Gpx4^KO^ (sg*Gpx4*) tumors (Fig. 2f, g). Efficient Fsp1 deletion was verified by immunohistochemistry (IHC), with less than 5% of tumors in the in sg*Fsp1* group still staining positive for Fsp1 (Extended Data Fig. 3a). We did not observe significant changes to the total number or size of tumors formed across the three biological groups (Extended Data Fig. 3b, c), which suggested that loss of Fsp1 or Gpx4 primarily impacts tumor progression rather than tumor initiation. We also observed that Fsp1 or Gpx4 deletion resulted in higher proportions of adenomas to adenocarcinomas compared to control tumors (Fig. 2h), indicating that Fsp1 and Gpx4 are functionally important for malignant disease progression. Additionally, there were no notable changes to tumor cell proliferation or apoptotic cell death (Extended Data Figs. 3d, e) in sg*Fsp1* or sg*Gpx4* tumors. In contrast, TUNEL staining of lung tumors demonstrated increased cell death with Fsp1 loss (Fig. 3l), suggesting that the decreased lung tumor burden with Fsp1 or Gpx4 genetic deletion is likely due to increased tumor cell ferroptosis. Perhaps most exciting though is the extent to which Fsp1 deletion phenocopied Gpx4 loss *in viv*o, as this phenotype, to our knowledge, has not yet been reported in any other cell lines, tissues, or disease models.

**Fig. 3:**
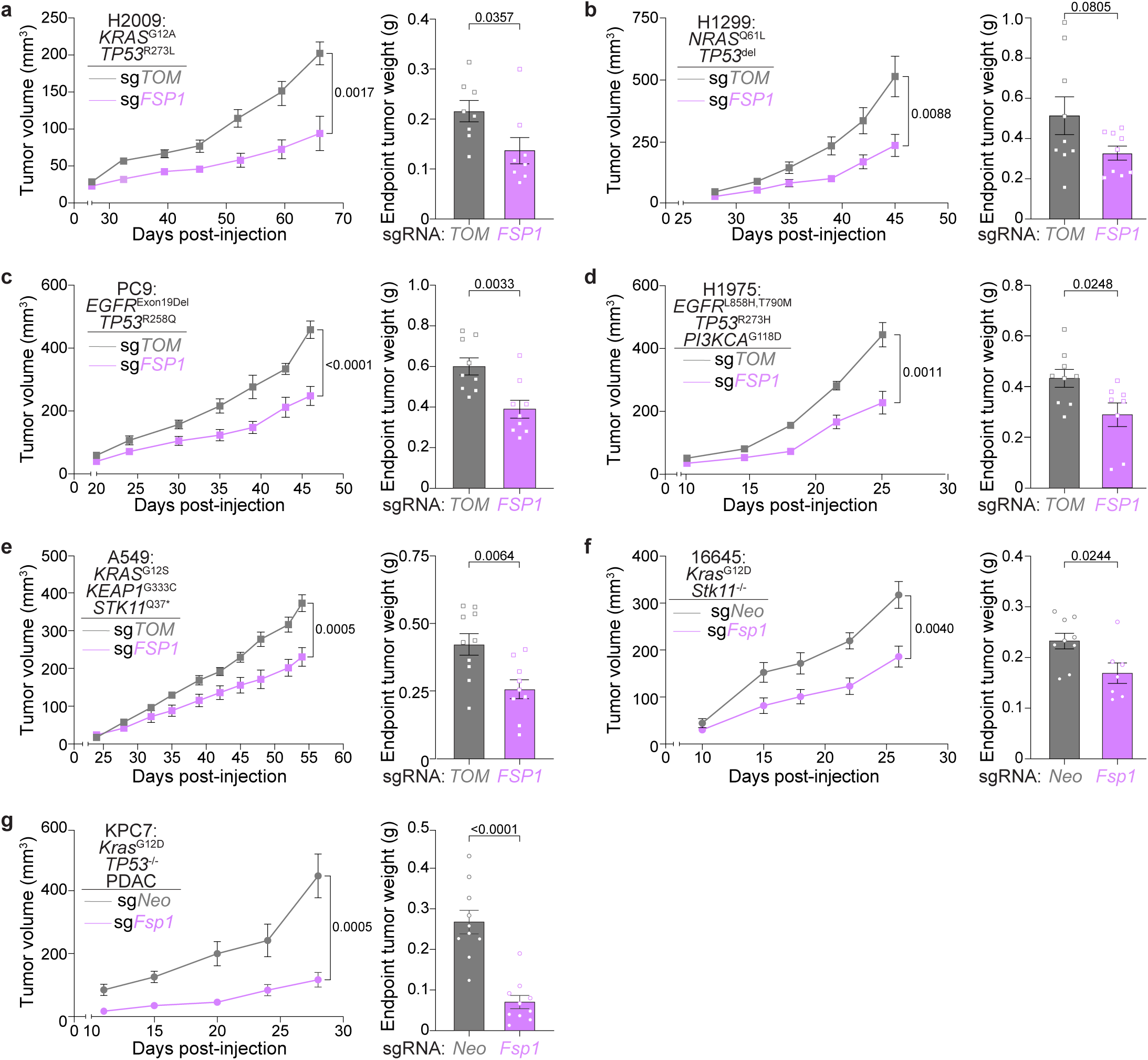
Sensitivity to FSP1 loss is not dependent on tumor mutations or lineage. Longitudinal tumor growth and endpoint tumor weights from indicated cell lines with CRISPR/Cas9-mediated knockout of FSP1 (sg*FSP1*) or control (sg*TOM*, sg*Neo*), transplanted as subcutaneous (subQ) xenograft tumors into NSG mice: **(a)** H2009, sg*FSP1* (n=8) or control (n=8). **(b)** H1299, sg*FSP1* (n=9) or control (n=9). **(c)** PC9, sg*FSP1* (n=9) or control (n=9). **(d)** H1975, sg*FSP1* (n=8) or control (n=9). **(e)** A549, sg*FSP1* (n=9) or control (n=10). **(f)** 16645, sg*Fsp1*(n=7) or control (n=9). **(g)** KPC7 (murine PDAC), sg*Fsp1* (n=9) or control (n=10). Data are represented as mean values, error bars represent SEM, significance determined via two-sided student’s t test (panels a-g)

### Fsp1 is required uniquely *in vivo*

The discovery of FSP1 as a key regulator of ferroptosis arose from genetic screens to identify ferroptosis suppressors in the absence of functional GPX4^8,9^. Thus, FSP1 is generally thought to control a secondary anti-ferroptotic axis whose function, at least *in vitro*, is masked when GPX4 is intact. Accordingly, we observed that CRISPR/Cas9-mediated Fsp1 deletion in KP LUAD cells had no impact on cell viability and clonogenicity *in vitro* (Fig. 2i, Extended Data Fig. 4a). As expected, Fsp1-knockout (Fsp1^KO^) cells were more sensitive to RSL3 treatment than isogenic Fsp1-wildtype (Fsp1^WT^) cells and protected from RSL3-killing by LIP1 (Fig. 2j). Despite the remarkable restriction of KP LUAD tumorigenesis by Fsp1 loss *in vivo*, our *in vitro* studies indicated in contrast that Fsp1’s anti-ferroptotic function was not required by KP LUAD cells in the presence of Gpx4.

We sought to dissect this intriguing dichotomy and better understand the distinct *in vivo* dependency of lung tumors on Fsp1. We first performed a series of syngeneic transplantation experiments using isogenic KP, Fsp1^KO^ and Fsp1^WT^ cell lines with no proliferation differences *in vitro* to systematically characterize the impact of Fsp1 loss on tumor growth and further interrogate whether Fsp1-mediated tumor differences were specific to *in vivo* growth conditions (Fig. 2i, Extended Data Fig. 4b). Consistently, we found that Fsp1^KO^ cells formed significantly smaller tumors in both immunocompetent subcutaneous xenografts (Extended Data Fig. 4c, d) as well as orthotopic lung tumors (Fig. 2k, Extended Data Fig. 4e, f). Robust suppression of Fsp1^KO^ orthotopic lung tumor growth was similarly observed in immunodeficient athymic (NU/J) mice (Extended Data Fig. 4g), as well as in an immunogenic mouse model^43^ (Extended Data Fig. 4h). We also observed the same phenotype in age- and sex-matched Fsp1 wildtype versus Fsp1 knockout mice (Extended Data Fig. 4i). These studies collectively provide strong evidence that Fsp1’s pro-tumorigenic effect is primarily cell-intrinsic. Fsp1 overexpression (Fsp1^OE^) in KP LUAD cells exhibited no *in vitro* growth advantage (Extended Data Fig. 4j) but had enhanced resistance to RSL3-killing (Extended Data Fig. 4k) and accelerated growth of xenograft tumors *in vivo* (Extended Data Fig. Fig. 4l, m). Altogether, these results suggest that Fsp1 is essential for LUAD cells’ survival *in vivo* and that increased Fsp1 expression is sufficient to promote *Kras-*mutant tumorigenesis *in vivo*.

**Fig. 4:**
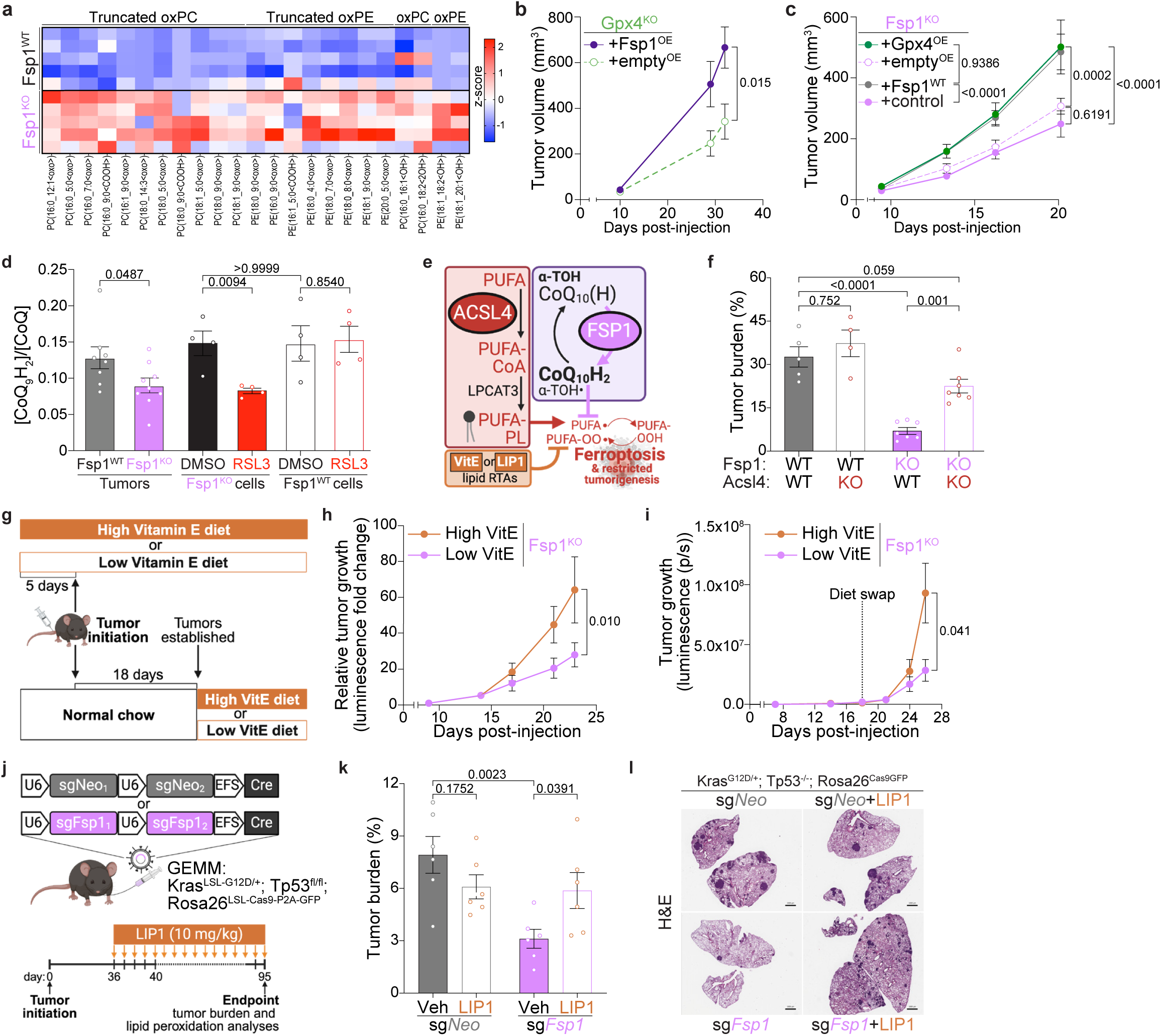
FSP1 is required for the suppression of ferroptosis *in vivo*. **(a)** Heatmap of indicated oxidized PE/PC lipid species detected via LC/MS from KP, Fsp1^KO^ and Fsp1^WT^ orthotopic lung tumors. **(b)** Longitudinal growth of KP, Gpx4^KO^ subQ xenograft tumors with Fsp1^OE^ (n=10) versus control (empty^OE^, n=9) in C57BL/6J mice. **(c)** Longitudinal growth of KP, Fsp1^KO^ subQ xenograft tumors with either Fsp1 restoration (Fsp1^WT^, n=9), Gpx4 overexpression (Gpx4^OE^, n=8), or controls (empty^OE^, n=9; Fsp1^KO^ n=8) in C57BL/6J mice. **(d)** Ratio of CoQ_9_H_2_/CoQ_9_ detected via LC/MS in KP, Fsp1^KO^ and Fsp1^WT^ orthotopic lung tumors versus cells treated with DMSO or RSL3 (0.5 µM) for 8 hours. **(e)** Schematic depicting Acsl4’s pro-ferroptotic function. **(f)** Tumor burden in C57BL/6J mice with KP, Fsp1^KO^ and Fsp1^WT^ orthotopic lung tumors with CRISPR/Cas9-mediated Acsl4 or control (Neo) deletion (WT n=5, Acsl4^KO^ n=4, Fsp1^KO^ n=7, Fsp1^KO^Acsl4^KO^ n=7). **(g)** Schematic depicting dietary Vitamin E (VitE) manipulation studies in (h) and (i). **(h)** Longitudinal lung tumor growth (measured via bioluminescence normalized to first timepoint) in C57BL/6J mice orthotopically transplanted with KP, Fsp1^KO^ cells and receiving high (n=8) or low (n=7) VitE diets *ad libitum* 5 days pre-tumor initiation. **(i)** Longitudinal lung tumor growth (measured via bioluminescence) in C57BL/6J mice orthotopically transplanted with KP, Fsp1^KO^ cells and placed on high (n=6) or low (n=6) VitE diets *ad libitum* after tumor establishment (day18). **(j)** Schematic of KP LUAD GEMMs intratracheally infected with pUSEC lentiviruses containing double sgRNAs targeting Neo (control, n=12) or Fsp1 (n=12). Mice were dosed with Liproxstatin-1 (LIP1, n=6 per genotype) or Vehicle (Veh, n=6 per genotype) daily starting 5 weeks post-tumor initiation. **(k)** Tumor burden of experiment in (j). **(l)** Representative H&E of tumors of experiment in (j). Scale bars: 100µm. Data are represented as mean values, error bars represent SEM, significance determined via two-sided student’s t test (panel b, d, f, h, i, k) or two-way ANOVA with Tukey’s multiple comparison test (panel c).

### FSP1 dependency is not mutation-specific

To more thoroughly interrogate the functional importance of Fsp1 for tumorigenesis and the translational potential for targeting Fsp1 in cancer, we assessed the requirement of FSP1 for growth of xenograft tumors from human LUAD cell lines harboring a variety of driver mutations, including *KRAS, NRAS, EGFR, TP53, STK11* and *KEAP1*. We found that CRISPR/Cas9-mediated FSP1 genetic deletion consistently and markedly decreased tumor growth in all models tested. Specifically, Fsp1^KO^ (sg*FSP1*) in H2009 (*KRAS*, *TP53* mutant), H1299 (*NRAS*, *TP53* mutant), PC9 and H1975 (*EGFR, TP53* mutant) LUAD cells led to robust suppression of subcutaneous tumor growth (Fig. 3a-d), but, analogous to aforementioned studies, did not impact proliferation or viability of cells *in vitro*, unless treated with RSL3 (Extended Data Fig. 5a-d). To assess whether *KRAS*-mutant tumors with *STK11* (LKB1) and/or *KEAP1* mutation also depend on FSP1, we performed CRISPR/Cas9-mediated FSP1 deletion in A549 (*KRAS, KEAP1, STK11* mutant) and 16645 (*Kras* mutant, *Stk11*-null) LUAD cells. Again, we observed a dependency for FSP1 *in vivo* (Fig. 3e, f) but not *in vitro* (Extended Data Figure 5e-f). Together these data provide further evidence that FSP1 is indeed required for *in vivo* LUAD tumor growth irrespective of driver and co-mutations.

Finally, we asked if Kras-driven tumors of a different lineage also exhibited a requirement for Fsp1 *in vivo*. Excitingly, transplantation of pancreatic adenocarcinoma (PDAC) *Kras*, *Tp53* mutant (KPC7) cells with CRISPR/Cas9-mediated Fsp1 deletion also led to a robust tumor suppression *in vivo* (Fig. 3g, Extended Data Fig. 5g), mirroring the phenotype observed consistently in LUAD models and suggesting that FSP1 dependency *in vivo* may extend to other tissue lineages.

### Ferroptosis inhibition rescues Fsp1 loss

We hypothesized that defective regulation of lipid peroxidation and subsequent induction of ferroptosis was the mechanism underlying FSP1^KO^ tumorigenic suppression. To assess this, we first performed epilipidomic analysis of Fsp1^WT^ and Fsp1^KO^ orthotopic lung tumors and observed that Fsp1^KO^ tumors exhibited increased abundance of oxidized and truncated PC/PE species, indicative of increased cellular lipid peroxidation (Fig. 4a). We then asked whether ectopic overexpression of Fsp1 or Gpx4 (Extended Data Fig. 6a, b) was sufficient to restore the growth of Gpx4 ^KO^ or Fsp1^KO^ tumors, which would shed light about the capacity for each ferroptosis defense arm to compensate for the loss of the other. Indeed, we found that Fsp1^OE^ and Gpx4^OE^ effectively restored the growth of Gpx4^KO^ and Fsp1^KO^ tumors, respectively (Fig. 4b-c, Extended Data Fig. 6c, d). These data implicated that tumors require extensive buffering capacity against lipid peroxidation and that while Gpx4 and Fsp1 act through distinct pathways, each can compensate for the loss of the other.

Mechanistically, Fsp1 has been shown to suppress ferroptosis by reducing CoQ, an endogenous RTA, and this function is dependent on its localization to the plasma membrane. We utilized a LC/MS-based, quantitative method to determine the abundance of CoQ in tumors, which revealed that Fsp1^KO^ tumors had a significantly decreased ratio of reduced CoQ_9_H_2_ to oxidized CoQ_9_ (Fig. 4d). This decreased ratio was not present in genotypically matched cells at baseline *in vitro*, but interestingly upon treatment with RSL3, Fsp1^KO^ cells mimicked the decreased CoQ_9_H_2_/CoQ_9_ observed in tumors. Prior studies have described that mutation of the myristoylation sequence that targets FSP1 to the plasma membrane renders FSP1 unable to protect against lipid peroxidation *in vitro*^8,9^, which we also observed in KP LUAD cells (Extended Data Fig. 6e, f). We anticipated that expression of this mutant Fsp1 (Fsp1^mut^) would be similarly unable to restore Fsp1^KO^ tumor growth. Indeed, while Fsp1^WT^-restored tumors grew quickly, both Fsp1^mut^ and Fsp1^KO^ tumors had decreased growth (Extended Data Fig. 6g). These data suggest that tumors require Fsp1 enzymatic activity and subcellular localization *in vivo* to specifically buffer against ferroptosis.

Next, we asked whether ferroptosis suppression, via several orthogonal approaches including tumor-specific Acsl4 knockout (Acsl4^KO^), dietary VitE supplementation, and LIP1 treatment would restore Fsp1^KO^ tumor growth. Loss or inhibition of Acsl4, which limits PUFA-PL supply and restricts lipid peroxidation (Fig. 4e), has previously been shown to be protective against ferroptosis in Gpx4^KO^ cells *in vitro*^17–19^. Therefore, we sought to determine whether genetic deletion of Acsl4 would suppress ferroptosis and rescue the growth of Fsp1^KO^ tumors *in vivo*. We performed CRISPR/Cas9-mediated genetic deletion of Acsl4 (sg*Acsl4*) in isogenic KP, Fsp1^KO^ and Fsp1^WT^ cells. We observed that loss of Acsl4 did not impact cell viability but robustly rescued RSL3-induced ferroptosis *in vitro*, in both Fsp1^WT^ and Fsp1^KO^ cells (Extended Data Fig. 6h). Importantly, Acsl4 deletion significantly restored Fsp1^KO^ lung tumor growth *in vivo* (Extended Data Fig. 6i) and mice with Acsl4, Fsp1-double knockout tumors were found to have increased endpoint disease burden (Fig. 4f, Extended Data Fig. 6j) and decreased overall survival (Extended Data Fig. 6k). Furthermore, tumor-specific Acsl4^KO^ led to a basal acceleration of tumor growth (Extended Data Fig. 6i) and decreased overall survival (Extended Data Fig. 6k), suggesting that decreased accumulation of PUFA-PL in lung tumors may also protect against ferroptosis *in vivo*.

VitE is nature’s premier lipophilic antioxidant and is specifically able to scavenge lipid radicals and protect against ferroptosis (Fig. 4e). Therefore, we decided to test whether dietary VitE supplementation could rescue the growth of Fsp1^KO^ lung tumors. High versus low VitE diets *ad libitum* was provided to animals either before or after tumor initiation (Fig. 4g), and we found that increased dietary VitE accelerated Fsp1^KO^ tumor growth (Fig. 4h, i), with no impact on growth of Fsp1^WT^ tumors (Extended Data Fig. 6l).

Lastly, since LIP1 suppressed tumor lipid peroxidation and entirely mitigated the impact of Gpx4 loss on KP LUAD tumorigenesis (Fig. 1b-d), we conducted parallel studies to investigate whether LIP1 also blunts ferroptosis in and restores Fsp1^KO^ tumorigenesis. In the orthotopic lung tumor model, we observed a tendency for Fsp1^KO^ tumors to grow more quickly with daily LIP1 administration (Extended Data Fig. 6m). In the KP LUAD GEMM (Extended Data Fig. 6n), which was conducted in accordance with experimental parameters for Gpx4^KO^ GEMM studies, LIP1 treatment significantly increased autochthonous Fsp1^KO^ (sg*Fsp1*) tumor burden (Extended Data Fig. 6o). Moreover, sg*Fsp1* tumors stained positively for 4-HNE, which was effectively suppressed by LIP1 treatment (Extended Data Fig. 6p). As observed with VitE supplementation, timing of LIP1 treatment did not appear to have a differential impact, as LIP1 treatment after tumors were already established was equally effective at restoring sg*Fsp1* lung tumorigenesis (Fig 4j-l). Altogether, these studies provide clear evidence that LUAD tumors require Fsp1 to protect against ferroptosis *in vivo* and suggest that increased lipid peroxidation is a barrier for tumor growth and progression.

### FSP1 inhibition extends overall survival

With the growing interest in harnessing ferroptosis to kill tumor cells, several FSP1 inhibitors have been developed^9,28^, though the majority are effective only *in vitro* against human FSP1 (hFSP1) in the context of GPX4 loss or inhibition (Extended Data Fig. 7a-d). Recently, icFSP1 was developed as the first human FSP1 (hFSP1) inhibitor with *in vivo* stability and efficacy, albeit solely in tumors with concomitant GPX4 loss^24^. Accordingly, we also saw that icFSP1 treatment *in vitro* did not impact cell viability (Extended Data Fig. 7e) unless RSL3 was added (Extended Data Fig. 7f). Given that murine and human FSP1 are highly conserved in sequence and structure, we generated a hybrid tumor model in which hFSP1 was expressed in murine KP, Fsp1^KO^ tumors. We observed that expression of either murine or human FSP1 resulted in similar acceleration of tumor growth compared to Fsp1^KO^ tumors (Extended Data Fig. 7g, h), which indicated that hFSP1 can functionally compensate for mFsp1 *in vivo*.

Using this hybrid tumor model, we next tested whether icFSP1 had therapeutic benefit against lung cancer. Excitingly, we found that FSP1 inhibition as a monotherapy improved overall survival of lung tumor-bearing mice, nearly to the extent of genetic Fsp1 deletion (Fig. 5a). To determine whether the therapeutic effect of icFSP1 was indeed due to the induction of ferroptosis, we asked whether LIP1, which we expected to inhibit lipid peroxidation induced by FSP1 inhibition, rescued tumor suppression in icFSP1-treated animals. In correlation with the survival data, we observed that icFSP1-treatment significantly decreased tumor growth when compared to vehicle treated animals, and importantly, concomitant LIP1 treatment abrogated the tumor-suppressive effect of icFSP1 treatment (Fig. 5b, Extended Data Fig. 7c). Further, to test whether icFSP1 was exerting an on-target, tumor-specific effect, we repeated the treatment study with an internal control— lung tumors expressing FSP1^Q319K^, a mutant that is resistant to icFSP1 but maintains anti-ferroptotic function (Extended Data Fig. 7a, b). We found that icFSP1 treatment specifically extended overall survival of mice bearing tumors with hFSP1^WT^ but not hFSP1^Q319K^ (Fig. 5c), indicating that indeed the tumor suppression seen with icFSP1 treatment was due to inhibitor activity directly on tumor cells.

**Fig. 5:**
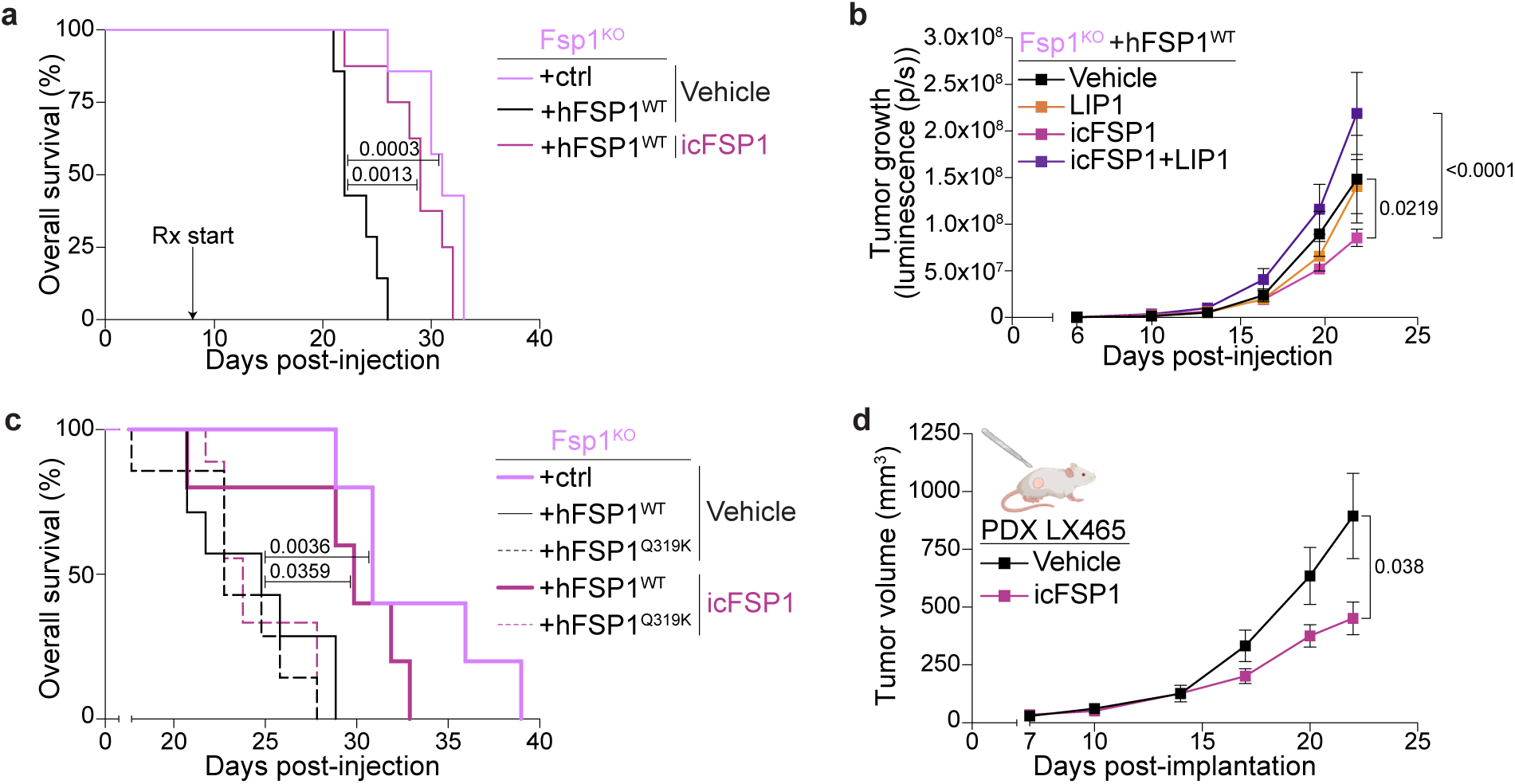
FSP1 is a viable therapeutic target for *KRAS*-mutant lung adenocarcinoma. **(a)** Overall survival of C57BL/6J mice with orthotopic lung tumors with Fsp1^KO^ (n=7) or hFSP1 re-expression, treated with either icFSP1 (n=8) or Vehicle (n=7). **(b)** Longitudinal lung tumor growth (measured via bioluminescence) in C57BL/6J mice orthotopically transplanted with hFSP1^WT^ cells and treated with either Vehicle (n=8), LIP1 (n=7), icFSP1 (n=8), or icFSP1+LIP1 (n=8). **(c)** Overall survival of C57BL/6J mice with Fsp1^KO^ (n=5), hFSP1^WT^, or hFSP1^Q319K^-expressing tumors, treated with icFSP1 (hFSP1^WT^, n=5; hFSP1^Q319K^, n=7) or Vehicle (hFSP1^WT^, n=7; hFSP1^Q319K^, n=7). **(d)** Longitudinal growth of PDX LX465 tumors treated with icFSP1 (n=10) or Vehicle (n=10) in NSG mice. Data are represented as mean values, error bars represent SEM, significance determined via two-way ANOVA with Tukey’s multiple comparisons test (panel b), two-sided student’s t-test (panel d), or Kaplan-Meier simple survival analysis (panels a, c).

We wondered whether icFSP1 treatment also impacted the tumor microenvironment (TME), but we found no differences in the proportion of total T cells, neutrophils, alveolar, or interstitial macrophages between the treatment groups (Extended Data Fig. 7j). These data further suggest that icFSP1 primarily exerts on-target, tumor-specific effects, though more granular immune profiling is needed to determine the potential impact, and thus potential side effects, of FSP1 inhibition on TME. Finally, we utilized a PDX model (LX465; *KRAS^G12D^, TP53*-mutant) to further assess the therapeutic potential of FSP1 inhibition. Excitingly, we found that icFSP1 treatment significantly decreased PDX tumor growth (Fig. 5d), further bolstering the translational potential of targeting FSP1 in LUAD patients. These pre-clinical studies with icFSP1 are the first to demonstrate that FSP1 is a promising therapeutic target, and our work highlight FSP1 inhibition as a novel therapeutic strategy for improving disease outcome in lung cancer patients.

## DISCUSSION

Cancer is a disease of hyperproliferation, and cancer cells increase their metabolic output to support sustained growth. This occurs at the cost of increased ROS production that can damage macromolecules, such as lipids, and have deleterious effects^44,45^. One such effect is excessive lipid peroxidation of membrane-associated PUFA-PLs, which can lead to ferroptosis, a unique non-apoptotic mechanism of cell death. Through our systematic interrogation of the key mechanisms that regulate aberrant lipid peroxidation *in vivo*, we demonstrated that ferroptosis constitutes a barrier to lung cancer and that lung tumors rely on Gpx4 and uniquely, Fsp1 to overcome ferroptosis and sustain tumorigenesis (Extended Data Fig. 7k, l).

Our work specifically highlights the dependence of lung tumors on Fsp1 *in vivo*, even in the presence of functional Gpx4 (Extended Data Fig. 7k). This is the first report of a context where Fsp1 and Gpx4 are independently required for ferroptosis suppression and therefore either can impact disease outcome. Moreover, analysis of GEMM and human lung tumors demonstrated that FSP1 levels increase during tumor progression and correlate with poor survival. In contrast, GPX4 levels are ubiquitous in tumors and most tissues. Given that germline Gpx4^KO^ mice are not viable^46^, whereas Fsp1^KO^ mice are with no notable physiological defects^20,47,48^, the therapeutic window for targeting FSP1 with fewer toxic side effects is expected to be much greater than for GPX4. Furthermore, loss of GPX4 is toxic to T cells which would suppress any anti-tumor immune responses^49,50^. Leveraging this unique requirement for Fsp1 in lung cancer, our studies demonstrate the therapeutic benefit of FSP1 inhibition in promoting ferroptosis and suppressing tumor growth and extending survival as a monotherapy in aggressive pre-clinical lung cancer models. Our studies underscore the value of *in vivo* pre-clinical models in capturing physiologically relevant metabolic dependencies that may be missed *in vitro* and pave the way for developing FSP1 inhibitors for clinical testing in lung cancer patients.

## Methods

### Cell lines

KP LUAD cell lines were obtained from the laboratory of Tyler Jacks. KP, Fsp1^KO^ cell lines were generated by transient transfection of PX458 (Addgene #48138) expressing an sgRNA targeting Fsp1. Single GFP positive clones were selected and Fsp1 loss was validated by western blot. The 16645 cell line was developed from Kras^G12D^ LKB1^−/−^ GEMM as previously described.^51^ KPC7 cells were obtained from the laboratory of Diane Simeone. All cell lines were maintained in DMEM or RPMI 160 (Corning) supplemented with 10% FBS (Sigma Aldrich) and gentamicin (Invitrogen) and were tested for mycoplasma regularly (PlasmoTest, InvivoGen). All mouse cell lines were authenticated by PCR genotyping. All human cell lines used were purchased from ATCC and were authenticated by short-tandem-repeat profiling. Genetic manipulation of cell lines was performed via lentiviral transduction of plasmids, detailed in next section, followed by either puromycin (7ug/mL) or hygromycin (800ug/mL) selection for 1 week.

### Cloning/lentivirus generation

CRISPR/Cas9-mediated knockout of target genes was achieved by cloning sgRNAs into pLenti-USEC or lentiCRISPRv2-puro vectors, as previously described^31^. In short, backbones were digested with Esp3I (New England Biosciences) and purified with a gel extraction kit (QIAGEN). sgRNAs were designed using CRISPick (Broad Institute), obtained from Integrated DNA Technologies (Coralville, IA), annealed, phosphorylated, Esp3I-digested, and ligated into the purified digested backbones using Quick Ligase (New England Biosciences). Double sgRNA ultramers were designed and generated as previously described^38^. Briefly, ultramers were Gibson assembled to digested pDonor_sU6 (Addgene #69351), Esp3I-digested, and ligated to purified digested backbones. Guide RNA sequences used to make gene knockouts can be found in Supplementary Data 4. Gpx4, mFsp1, and hFSP1 expression plasmids were generated using Gibson assembly of the respective cDNA into pLenti-v2-filler.

Lentivirus was generated by co-transfection of HEK293 cells with a viral vector and packaging plasmids psPAX2 (Addgene #12260) and pMD2.G (Addgene #12259) using PEI transfection reagent. Cell supernatant containing lentivirus was collected 72 hours after transfection and filtered through 0.45 µM PVDF filters. For *in vivo* experiments, lentivirus was concentrated by ultracentrifugation at 25,000rpm for 2 hours at 4°C. The viral pellet was resuspended in PBS and stored at −80°C until use. Viral titer was quantified with the use of a Cre-dependent GreenGo reporter cell line. For *in vitro* experiments, media containing virus was collected, filtered, and added directly to recipient cells with polybrene at 8 µg/mL for 48 hours before selection.

### Mouse models

All mouse experiments described in this study were approved by the NYU Institutional Animal Care and Use Committee (IACUC). Animals were housed according to IACUC guidelines in ventilated caging in a specific pathogen free animal facility. For all mouse studies, ≥ 4 mice were used for each experimental condition. *Kras*^LSL-G12D/+^; *Tp53*^fl/fl^; *Rosa26*^LSL-Cas^^9/LSL-Cas9^ (KPC GEMMs) were bred as previously described^31–35^. C57BL/6J (JAX strain #000664) mice with the appropriate genotype, aged 8 to 12 weeks were randomly selected to begin tumor initiation studies with pUSEC lentivirus. Care was taken to ensure each experimental arm had an equal number of male and female mice. Mice were intratracheally infected with lentiviruses as described and monitored until experimental endpoint. Tumor burden was quantified by hematoxylin and eosin (H&E) staining and analyzed using QuPath software as a measurement of total tumor area/total lung lobe area. All quantifications were done with investigator blinded to the respective sample genotypes. All transplantation experiments were performed using nude (JAX strain #002019), NOD SCID Gamma (NSG; JAX strain #005557F), C57BL/6J Fsp1-knockout (Conrad group^16^) or C57BL/6J wildtype (JAX strain #000664) mice aged 8 to 12 weeks old. For murine cell xenograft experiments, 100,000 cells in 100µl of phosphate-buffered saline (PBS) were injected subcutaneously into each flank of the mouse. For the xenograft studies in Figure 3 the number of cells injected per flank and whether they were injected with 50:50 PBS and Matrigel (Corning) are indicated as follows: H2009 (2 million cells +Matrigel), H1299 (1 million cells), PC9 (1 million cells), H1975 (2 million cells +Matrigel), A549 (1 million cells), 16645 (500,000 cells), KPC7 (250,000 cells). All human cell line xenograft experiments were carried out in male NSG mice unless specified. For the PDX experiment, tumors were implanted subcutaneously in male NSG mice as previously described^34^. Tumors were measured by caliper, and volume was calculated based on 0.5 x length x width^2^. The maximum tumor diameter permitted by our IACUC protocol was 2cm, and this was not exceeded in any experiment. For orthotopic lung tumor experiments, 100,000 luciferase-expressing cells in 200μl of PBS were injected intravenously into tail vein of male mice unless specified in the legend. Tumor growth was measured by bioluminescence (PerkinElmer IVIS Spectrum In Vivo Imaging System; D-luciferin, PerkinElmer #122799). Data were analyzed using Living Image software.

### Antioxidant/drug treatments

For LIP1 treatment, mice were dosed with 10 mg/kg Liproxstatin-1 (BOC Sciences) or Vehicle (2% DMSO + 40%PEG300 + 2%Tween80 in sterile H_2_O) by intraperitoneal injection for frequency and duration indicated in figure schematics. For icFSP1 treatment, mice were dosed with 50mg/kg icFSP1 (WuXi LabNetwork) or vehicle (45% PEG300 in sterile PBS) by intraperitoneal injection twice daily. High (TD.2108412) and low (TD.210841) irradiated Vitamin E diets were obtained from Inotivco and provided *ad libitum* for length of time indicated in figure legends. In all experiments, mice were randomly assigned to treatment group.

### Cell clonogenic and viability assays

Cell clonogenic assays were conducted by seeding 2000 cells per well into 12-well dishes (BD/Falcon) in RPMI-1640 media. After 12-16hrs, media containing RSL3 and/or LIP1 were added to wells. After 5 days of growth, plates were washed twice with PBS and stained with 0.5% crystal violet (Fisher Scientific) solution in 20% methanol. Plates were dried, scanned, and crystal violet was quantified by solubilization with 10% acetic acid and measurement of absorbance at 592nm by spectrometer (Molecular Devices). Cell viability assays were conducted by seeding 2000 cells per well into white-walled, clear-bottom, 96-well plates (Corning) in RPMI-1640 media. After 12-16hrs, media containing RSL3, LIP1, and/or FSEN1, iFSP1, or icFSP1 were added. After 3 days, cell viability was assessed by CellTiter-Glo (Promega) and luminescence was measured by spectrometer (Molecular Devices).

### Immunoblotting

Cells were plated to 75% confluency in a 6-well dish, and the following day cells were lysed on ice with Pierce RIPA buffer (ThermoScientific) containing 1x protease/phosphatase inhibitor cocktail (Thermo Fisher Scientific). Samples were sonicated in cooled 4°C water (12 rounds, 30 seconds on and 30 seconds off) and then centrifuged at 14000 rpm at 4°C for 15 minutes. The supernatant was collected, and protein was quantified using the DC Rad Protein Assay kit. Protein was diluted to 1 ug/uL with water and 4x NuPage LDS sample buffer, then boiled at 95°C for 10 minutes. 20 ug protein per well was loaded into Invitrogen 4-12% Bis-Tris gels and then transferred onto PVDF membranes using a standard protocol. PVDF membranes were then blocked using 5% BSA in TBST for 60 minutes at room temperature and incubated with primary antibodies in 5% BSA overnight at 4°C. Primary antibodies were obtained as follows: Gpx4 (Abcam), Fsp1 (Proteintech), Acsl4 (Santa Cruz Biotechnologies), pERK (Cell Signaling), ERK (Cell Signaling), and Hsp90 (BD Bioscences). The following day, membranes were washed in TBST and incubated with HRP-conjugated secondary antibodies for 1 hour at room temperature. Enhanced chemiluminescent horseradish peroxidase substrate (ThermoScientific SuperSignal West PICO Plus) was added to the membrane for 1 minute, and the resulting membrane was imaged using the General Electric Amersham Imager 680. For gel source data, see Supplementary Data 1.

### Oxidized lipidomics

For the Gpx4 KO experiment, cells were plated to 75% confluency in 6-well dish. The following day, cells were treated with DMSO, RSL3 (0.5uM), or RSL3(0.5µM) + LIP1(100nM). After 8hrs, cells were collected and washed in an antioxidant solution (PBS containing dibutylhydroxytoluene (BHT; 100 µM) and diethylenetriamine pentaacetate (DTPA; 100 µM)) and centrifuged. Supernatent was discarded and cell pellets were immediately frozen in liquid nitrogen and stored at –80 °C. Frozen samples were sent on dry ice to Wayne State Lipidomics Core for metabolite extraction and LC-MS analysis.

Fsp1 WT and KO tumor and cell lipidomic analysis were performed in the lab of Pyo Kang. Fsp1 WT and KO cells were plated at 75% confluency in 10 cm plates and treated as stated above. After euthanasia, animals were perfused with the antioxidant solution (+3.8% trisodium citrate), orthotopic tumors were microdissected, frozen in liquid nitrogen and stored at –80 °C. Mouse lung cancer tissues were cryopulverized and extracted with chloroform:methanol (2:1, v/v) at a tissue concentration of 25 mg mL⁻¹. For the cells, 1.0 × 10^7^ cells were extracted by one milliliter of chloroform:methanol (2:1, v/v). EquiSPLASH LIPIDOMIX® internal standard (Avanti Polar Lipids) was added to the extraction solvent at 0.1 µg mL⁻¹ per lipid class. Samples were sonicated on ice for 1 min using a VCX 130 probe sonicator (5 s on/off cycles), incubated for 30 min, and centrifuged at 17,000 × g for 20 min at 4 °C. The supernatant was was dried under vacuum (EZ-2 Elite, Genevac), reconstituted in 10% of the original volume with isopropanol, and transferred to glass autosampler vials for LC–MS analysis. Chromatographic separation was performed using a Waters ACQUITY UPLC CSH C18 column (100 × 2.1 mm, 1.7 µm) with a VanGuard precolumn (5 × 2.1 mm, 1.7 µm). Mobile phase A consisted of acetonitrile/water (60:40, v/v), and mobile phase B of isopropanol/acetonitrile (90:10, v/v), both containing 5 mM ammonium formate and 0.1% formic acid. The column was maintained at 65 °C with a flow rate of 0.6 mL min⁻¹. The gradient was as follows: 0–2 min, 15–30% B; 2–2.5 min, 30–48% B; 2.5–11 min, 48–82% B; 11–11.5 min, 82–99% B; 11.5–12 min, 99% B; 12–12.1 min, 99–15% B; and 12.1–16 min, hold at 15% B for re-equilibration. Mass spectrometry was performed on a Q Exactive Plus Quadrupole-Orbitrap (Thermo Fisher Scientific) equipped with a heated electrospray ionization (HESI) source and operated in negative ion mode. Source settings were: sheath gas, 60 a.u.; auxiliary gas, 25 a.u.; sweep gas, 2 a.u.; spray voltage, 3.0 kV; capillary temperature, 320 °C; S-lens RF level, 50%; and auxiliary gas heater temperature, 370 °C. Parallel reaction monitoring (PRM) was carried out with the following parameters: resolution, 17,500 at m/z 200; AGC target, 2 × 10⁵; maximum injection time, 50 ms; isolation window, m/z 1.2; and stepped normalized collision energies of 20, 30, and 40. The oxidized lipid analysis was adapted by previous PRM based analysis^52^. To generate the PRM inclusion list, pooled sample extracts were first analyzed in DDA mode to identify the most abundant polyunsaturated PC and PE species using Lipostar2^53^. These precursors were subjected to in silico oxidation using LPPtiger2 to predict candidate oxidized lipids. A semi-targeted DDA experiment was then conducted to confirm precursor detectability and finalize the inclusion list. ^52^ Data were analyzed using Skyline (v24.1)^54^. Quantification was based on fragment anions derived from oxidized fatty acyl chains, with peak areas normalized to PC(15:0/18:1(d7)) or PE(15:0/18:1(d7)) internal standards from the EquiSPLASH LIPIDOMIX® mixture.

### CoQ_9_ and CoQ_9_H_2_ analysis

CoQ_9_ and CoQ_9_H_2_ analysis was conducted by modification of previous study^55^. The D_9_-CoQ_10_ standard was purchased from IsoSciences (Ambler, PA, USA). The D_6_-CoQ_10_H_2_ standard was synthesized from D_6_-CoQ_10_, which was obtained from Good Laboratory Practice Bioscience (Montclair, CA, USA). The CoQ_9_ standard was purchased from Tokyo Chemical Industry (Tokyo, Japan), and CoQ_9_H_2_ was synthesized from CoQ_9_. D_6_-CoQ_10_H_2_ and D_9_-CoQ_10_ were used as internal standards for the CoQ_9_H_2_ and CoQ_9_, respectively. Chloroform: methanol (2:1, v/v), containing internal standards of 0.5 µM of D_6_-CoQ_10_H_2_ and D_6_-CoQ_10_ was used to extract mouse lung cancer tissue at a tissue concentration of 25 mg mL⁻¹. For the cells, 1.0 × 10^7^ cells were extracted by one milliliter of same extraction solvent. Samples were then sonicated on ice for 1 min using a VCX 130 probe sonicator (5 s on/off cycles), incubated for 30 min, and centrifuged at 17,000 × g for 20 min at 4 °C. A 100 µL aliquot of the supernatant was transferred to glass autosampler vials for LC–MS/MS analysis. The liquid chromatography conditions were identical to those used for oxidized lipid analysis. Q Exactive Plus MS was operated in positive ion mode. Source settings were as follows: sheath gas, 60 a.u.; auxiliary gas, 25 a.u.; sweep gas, 2 a.u.; spray voltage, 3.0 kV; capillary temperature, 320 °C; S-lens RF level, 50%; and auxiliary gas heater temperature, 370 °C. The mass range was m/z 120–1,200; resolution, 70,000 at m/z 200; AGC target, 1 × 10^6^; and maximum injection time, 100ms. By using EL–MAVEN (v0.12.0), the LC-MS peaks of CoQ_9_, CoQ_9_H_2_, D_6_-CoQ_10_H_2_, D_6_-CoQ_10_, and D_9_-CoQ_10_ were identified by matching with standard library and their peak areas were extracted with 10 ppm error range. Standard curves were generated using known concentrations of CoQ_9_H_2_ and CoQ_9_, and used to calculate the concentrations of CoQ_9_H_2_ and CoQ_9_ in the samples fallowing an algorithm described in a previous study^55^.

### Immunohistochemistry

Tumor bearing mice were euthanized by carbon dioxide asphyxiation, after which the lungs were dissected and fixed in 4% PFA solution overnight. Fixed lungs were washed with PBS 2x, transferred, and stored in 70% ethanol, until cassette loading and paraffin embedding. Sections were cut and stained with H&E. For immunohistochemistry with the exception of TUNEL staining, sections were immunostained on a Leica BondRX automated stainer according to the manufacturer’s instructions. In brief, tissues underwent deparaffinization online, followed by epitope retrieval for 20 minutes at 100° with Leica Biosystems ER2 solution (pH9, cat#AR9640), endogenous peroxidase activity blocking with H2O2, and non-specific binding site blocking with Rodent Block M (Biocare, cat# RBM961L) and Bond Primary Antibody Diluent (Leica Biosystems, cat# AR9352). Sections were then incubated with primary antibodies against Gpx4 (Abcam), Fsp1 (courtesy of Dr. Marcus Conrad), 4-HNE (JaICA), Ki67, and Ccasp3 for 60 minutes at room temperature. Primary antibodies were detected with anti-rat HRP-conjugated polymer (Biocare, cat# BRR4016H), 3,3’-diaminobenzidine (DAB) substrate (provided in the Leica BOND Polymer Refine Detection System, cat# DS9800), and for 4-HNE staining Bond DAB Enhancer (Leica Biosystems, cat# AR9432). Following counter-staining with hematoxylin, slides were scanned at 40X on a Hamamatzu Nanozoomer (2.0HT). TUNEL staining was performed by Histowiz Inc. (histowiz.com) according to their protocol.

For OPAL imaging, coronal 5 µm sections were immunostained on a Leica BondRx auto-stainer according to the manufacturer’s instructions. In brief, sections were deparaffiinized online and then treated with 3% H_2_O_2_ to inhibit endogenous peroxidases, followed by antigen retrieval with either ER1 (Leica, AR9961; pH6) or ER2 (Leica, AR9640; pH9) retrieval buffer at 100o for 20 minutes. After blocking with either Rodent Block M (Biocare, RBM961L) or Primary Antibody Diluent (Leica, AR93520), slides were incubated with the first primary antibody (FSP1, courtesy of Marcus Conrad, pERK1/2, CST, GPX4, Abcam) and secondary HRP polymer pair, followed by HRP-mediated tyramide signal amplification with a specific Opal® fluorophore. Once the Opal® fluorophore was covalently linked to the antigen, primary and secondary antibodies were removed with a heat retrieval step. This sequence was repeated three more times with subsequent primary and secondary antibody pairs, using a different Opal fluorophore with each primary antibody (see table below for primary antibody sequence and reagent details). After antibody staining, sections were counterstained with spectral DAPI (Akoya Biosciences, FP1490) and mounted with ProLong Gold Antifade (ThermoFisher Scientific, P36935). Semi-automated image acquisition was performed on an Akoya Vectra Polaris (PhenoImagerHT) multispectral imaging system. Slides were scanned at 20X magnification using PhenoImagerHT 2.0 software in conjunction with Phenochart 2.0 and InForm 3.0 to generate unmixed whole slide qptiff scans. All image files were uploaded to the NYUGSoM’s OMERO Plus image data management system (Glencoe Software).

### Flow Cytometry

Lung tissue was processed into a single cell suspension for flow cytometry as previously described^43,56^. Briefly, prior to euthanasia, mice were injected with 2 ug of Anti-mouse CD45-APC conjugated antibody (Biolegend, Clone 30-F11, 103111) retro-orbitally. Lungs were harvested and digested with Collagenase (Sigma-Aldrich, C5138) and deoxyribonuclease I (Sigma-Aldrich, DN25) followed by red blood cell lysis. Single cells were then resuspended in FACS buffer and stained using the following antibodies: CD45 (Biolegend, 103132), CD11b (Biolegend, 101216), CD11c (Biolegend, 117324), Ly6G (Biolegend, 127622), MHCII (BD, 748708), CD103 (Biolegend, 121433), CD64 (Biolegend, 139309), SiglecF (BD, 740956), MertK (R&D, BAF591), CD45 (BD, 748371), CD3e (BD, 740854), CD4 (Invitrogen, MCD0428), CD8a (EBioscience, 563152), Secondary (Streptavidin) (BD, 564176). Samples were run on the BD LSRFortessa and analyzed on FlowJo version 10. Gating strategy can be accessed in Supplementary Data 2.

### TCGA analyses

Gene expression profiles of primary tumors and relevant clinical data of 515 LUAD patients were obtained from The Cancer Genome Atlas6 (TCGute.org). GPX4 and FSP1 (AIFM2) mutational status of TCGA tumor samples was retrieved from cBioPortal151 using the TCGA PanCancer Atlas collection (gdc.cancer.gov/about-data/publications/pancanatlas). For survival data, patients were stratified based on GPX4 or FSP1 expression and overall survival rates were plotted to compare high-GPX4/FSP1 expressing patients (top 50% above median expression) with the rest of the cohort (n=464 patients). All survival analyses were conducted using the survival curve analyses in GraphPad Prism v9.

### Statistics and Reproducibility

Statistical analysis was performed using GraphPad Prism v9. All data is expressed as mean plus standard error of the mean, unless otherwise specified. Data was analyzed by statistical test indicated in figure legends. All tests were two-tailed and replicates are biological unless otherwise stated. All western blots were replicated at least 3 times with results reproducible of data shown in figures. All *in vitro* assays were replicated at least 3 times with a minimum of n=3 biological replicates per group for statistical power. For all *in vivo* experiments, the minimum sample size was 4 independent mice/tumors and respective sample size per genotype/condition is further specified in the figure legend. Sample size was not calculated but was chosen in each experiment based on previous experience with various models and to ensure that there were enough samples for statistical power. All *in vivo* experiments were replicated at least twice with results reproducible of data shown in figures. When representative images are shown, a minimum of 3 samples from the larger cohort were stained from each group. In the case of representative MRI images, all animals from the cohort were imaged. During sample processing and analysis for the lipidomic the samples were given numeric IDs which after analysis, were unblinded and graphed. For histological and immunohistochemistry analysis, researcher was blinded to the sample condition. The investigators were not blinded during most other data collection or analysis. Biorender^57^ was used to make the schematics and diagrams.

## Data Availability

All data and raw gel images are included with the paper. Raw lipidomic data have been deposited in the MassIVE data base (https://massive.ucsd.edu/) under accession number MSV000098883. Analyzed lipidomic data are available in Supplementary Data 3. Gene expression profiles of primary tumors and relevant clinical data of 515 LUAD patients were obtained from The Cancer Genome Atlas6 (TCGute.org). GPX4, FSP1 (AIFM2), KRAS, EGFR, KEAP1, and STK11 mutational status of TCGA tumor samples was retrieved from cBioPortal151 using the TCGA PanCancer Atlas collection (gdc.cancer.gov/about-data/publications/pancanatlas). All other materials are available upon request from the corresponding author (T.P.). Source data are provided with this paper.

## Acknowledgements

We dedicate this work to Judy Teixeira, whose love, strength, and spirit continue to inspire us. We thank Gyles Ward and Cynthia Loomis from the Experimental Pathology Core [RRID:SCR_017928] at NYU Langone Health, which is partially supported by the Cancer Center Support Grant P30CA016087 at NYU Langone’s Laura and Isaac Perlmutter Cancer Center, for support with IHC staining and imaging. The PhenoImagerHT Mutlispectral imaging system was initially purchased through a Shared Instrumentation Grant S10OD021747. We thank the Wayne State University School of Medicine Lipidomics Core Facility, which is supported in part by National Center for Research Resources, National Institutes of Health Grants S10RR027926 and S10OD032292, for support with LC-MS. We also recognize that the analysis of Fsp1 oxidized lipids and CoQ_9_ were conducted by the Proteomics and Metabolomics Core at the College of Pharmacy, Seoul National University, South Korea. We thank R. Possemato and S. Kotschi for critical reading of this manuscript. T. P. is supported by NIH grants (R37CA222504, R01CA227649, R01CA283049, R01CA262562) and an American Cancer Society Research Scholar Grant (RSG-17-20001–TBE). K.W. is supported by the Ruth L. Kirschstein Individual Predoctoral NRSA fellowship (F30CA275258) and NIH training grants (T32GM136573, T32GM136542). A.J.V. is supported by the NIH training grant (T32GM136542). M.C. received funding from the Deutsche Forschungsgemeinschaft (DFG) (CO 291/7-1, the Priority Program SPP 2306 [CO 291/9-1, #461385412; CO 291/10-1, #461507177] and the European Research Council (ERC) under the European Union’s Horizon 2020 research and innovation program (grant agreement No. GA 884754).

## Author contributions

T.P., K.W., A.J.V. designed and directed the study. K.W., A.J.V., J.P.B., A.Z., M.N.N., M.M., and S.C. performed various *in vitro* and *in vivo* experiments reported in the study. R.P. performed FACS. Y.H. performed TCGA patient data analyses. S.M.K. performed Fsp1 epilipidomics and COQ measurement, T.H.K assisted with data analysis, and was supervised by Y.P.K. S.C. performed unbiased tumor burden quantification. S.R. maintained and genotyped all experimental mice. M.C., T.N., D.S., M.H., provided conceptual advice. T.P., K.W., and A.J.V. wrote the manuscript with comments and input from all authors.

## Extended Data Figure Legends

**Extended Data Fig. 1:**
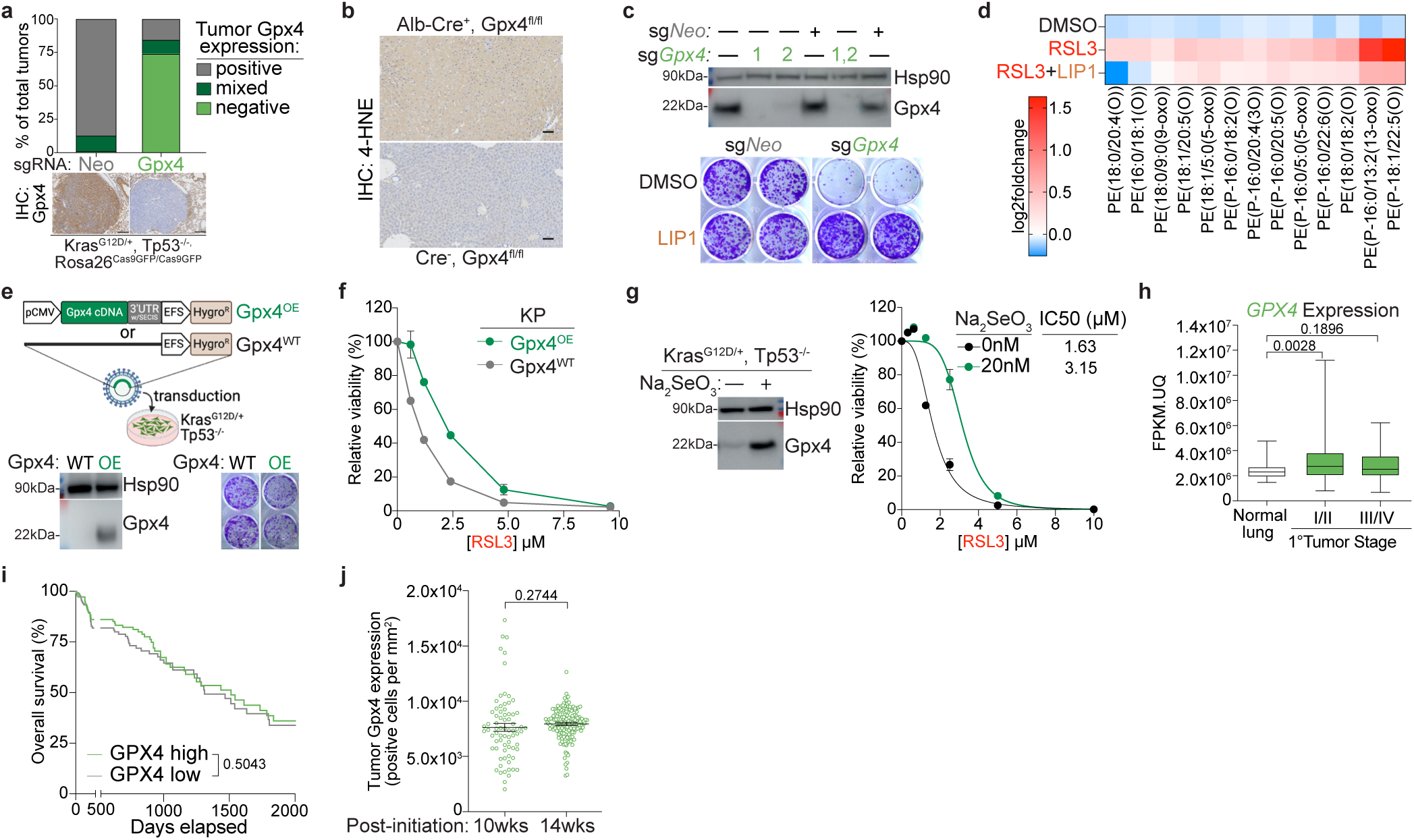
Gpx4 is required by lung cancer cells. **(a)** Quantification and representative images of Gpx4 IHC in KP LUAD GEMM tumors with knockout of either control (Neo, n=11) or Gpx4 (n=12). Scale bars: 200µm. **(b)** Representative 4-hydroxy-2-noneal (4-HNE) IHC of liver tissue from conditional *Gpx4*-knockout mice. Scale bars: 100µm. **(c)** Top: Western blot of KP LUAD cells with CRISPR/Cas9-mediated genetic deletion of Gpx4 with either two individual or duplexed sgRNAs. Bottom: Representative images of crystal violet clonogenic assay of KP LUAD cells with knockout of either control (Neo) or Gpx4. Cells were treated with 100nM LIP1. **(d)** Heatmap of LC-MS detection of oxidized phospholipids in KP LUAD cells treated with DMSO control, RSL3 (0.5µM), and RSL3 (0.5µM) + LIP1 (100nM) for 8 hours. **(e)** Schematic of Gpx4 ectopic overexpression (OE) method in KP LUAD cells. Western blot and representative images of crystal violet clonogenic assay of KP LUAD cells with wildtype (WT) or OE of Gpx4. **(f)** CellTiter-Glo Luminescence viability assay of KP, Gpx4^WT^ or Gpx4^OE^ cells upon increasing concentrations of RSL3 (n=5 per group). **(g)** Western blot of KP LUAD cells treated with 20nM Na_2_SeO_3_ or DMSO. CellTiter-Glo Luminescence viability assay of KP LUAD cells treated with 20nM Na_2_SeO_3_ or DMSO with increasing RSL3 addition (n=5 per group). **(h)** *GPX4* expression in *KRAS*-mutant primary LUAD tumors from TCGA, divided into early and late tumor stages (normal lung, n=54; stage I/II, n=354; stage III/IV, n=98). **(i)** Overall survival of *KRAS*-mutant LUAD patients (n=464) from TCGA, stratified by high vs low tumor *GPX4* expression. **(j)** Tumor Gpx4 IHC quantification of KP LUAD GEMM tumors at 10 weeks (n=4) vs 14 weeks (n=4) post-tumor initiation. Box plots indicate median (middle line), 25th, 75th percentile (box) and 5th and 95th percentile (whiskers). Data are represented as mean values, error bars represent SEM, significance determined via one-way ANOVA with multiple comparisons (panel h), two-sided student’s t-test (panel j) or Kaplan-Meier simple survival analysis (panel i). For gel source data, see Supplementary Data 1.

**Extended Data Fig. 2:**
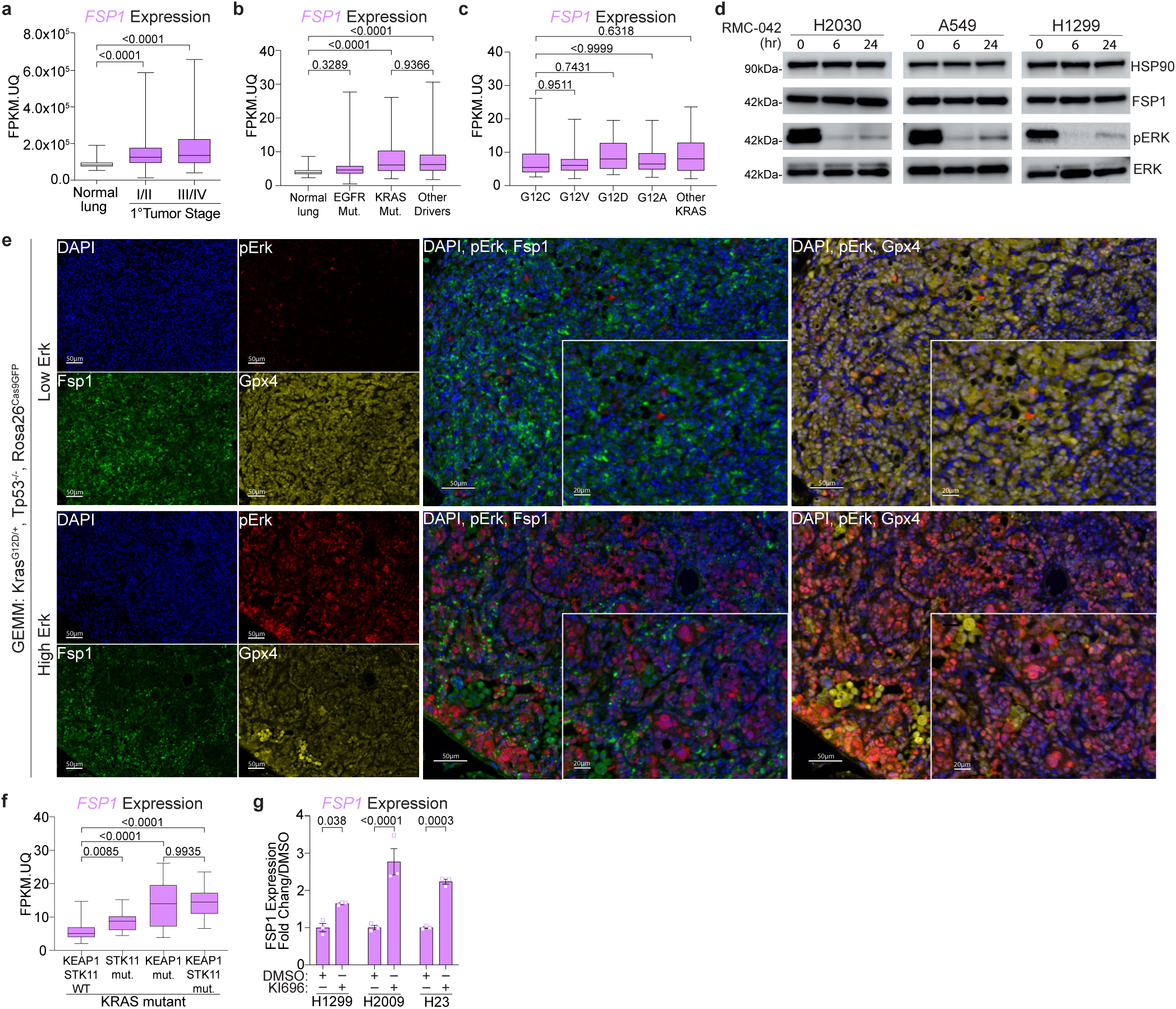
FSP1’s anti-ferroptotic function is not dependent on oncogenic signaling. **(a)** *FSP1* (*AIFM2*) expression of *KRAS*-mutant primary LUAD tumors from TCGA, divided into early and late tumor stages (normal lung, n=54; stage I/II, n=354; III/IV, n=98). **(b)** *FSP1* (*AIFM2*) expression of *KRAS*-mutant primary LUAD tumors from TCGA, separated by the *KRAS* mutation (G12C n=51; G12V n=32; G12D n=17; G12A n=16; other n=20). **(c)** *FSP1* (*AIFM2*) expression of primary LUAD tumors from TCGA, separated by oncogenic driver mutation (normal lung, n=59; EGFR, n=71; KRAS, n=135; other n=307). **(d)** Western blot of KP LUAD human cell lines treated with 50nM RMC-042 for indicated durations. **(e)** Representative multi-IF images of KP tumors for markers indicated. Panels are 10X, scale bars: 50µm; insets are 20X, scale bars: 20µm. **(f)** *FSP1* (*AIFM2*) expression of *KRAS*-mutant primary LUAD tumors from TCGA, separated by tumor co-mutation status (*KEAP1*/*STK11* WT n=88; *STK11* mutant n=20; *KEAP1* mutant n=16; *KEAP1/STK11* mutant n=11). **(g)** *FSP1* (*AIFM2*) expression in KP LUAD human cell lines treated with an Nrf2 activator, KI696, for 5 days (n=3 per group). Box plots indicate median (middle line), 25th, 75th percentile (box) and 5th and 95th percentile (whiskers). Data are represented as mean values, error bars represent SEM, significance determined via one-way ANOVA with multiple comparisons (panels a, b, c, f, g). For gel source data, see Supplementary Data 1.

**Extended Data Fig. 3:**
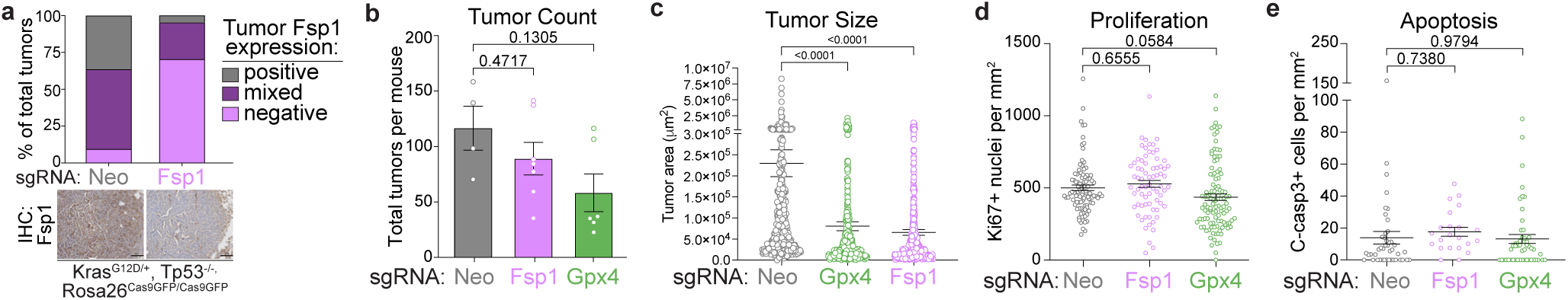
Neither Fsp1 nor Gpx4 have an impact on tumor initiation, proliferation, or apoptosis. **(a)** Quantification and representative images of tumor Fsp1 IHC in KP LUAD GEMMs with knockout of either control (Neo, n=11) or Fsp1 (n=14). Scale bars: 100µm. **(b)** Tumor number quantification in KP LUAD GEMMs with tumor-specific knockout of either control (Neo, n=4), Fsp1 (n=7), or Gpx4 (n=6). **(c)** Individual area of KP LUAD GEMM tumors with knockout of either control (Neo, n=465), Fsp1 (n=453), or Gpx4 (n=384). **(d)** Quantification of Ki67 IHC of KP LUAD GEMM tumors with knockout of either control (Neo, n=355), Fsp1 (n=268), or Gpx4 (n=291). **(e)** Quantification of cleaved caspase-3 IHC of KP LUAD GEMM tumors with knockout of either control (Neo, n=46), Fsp1 (n=22), or Gpx4 (n=52). Data are represented as mean values, error bars represent SEM, significance determined via one-way ANOVA with multiple comparisons (panels b-e).

**Extended Data Fig. 4:**
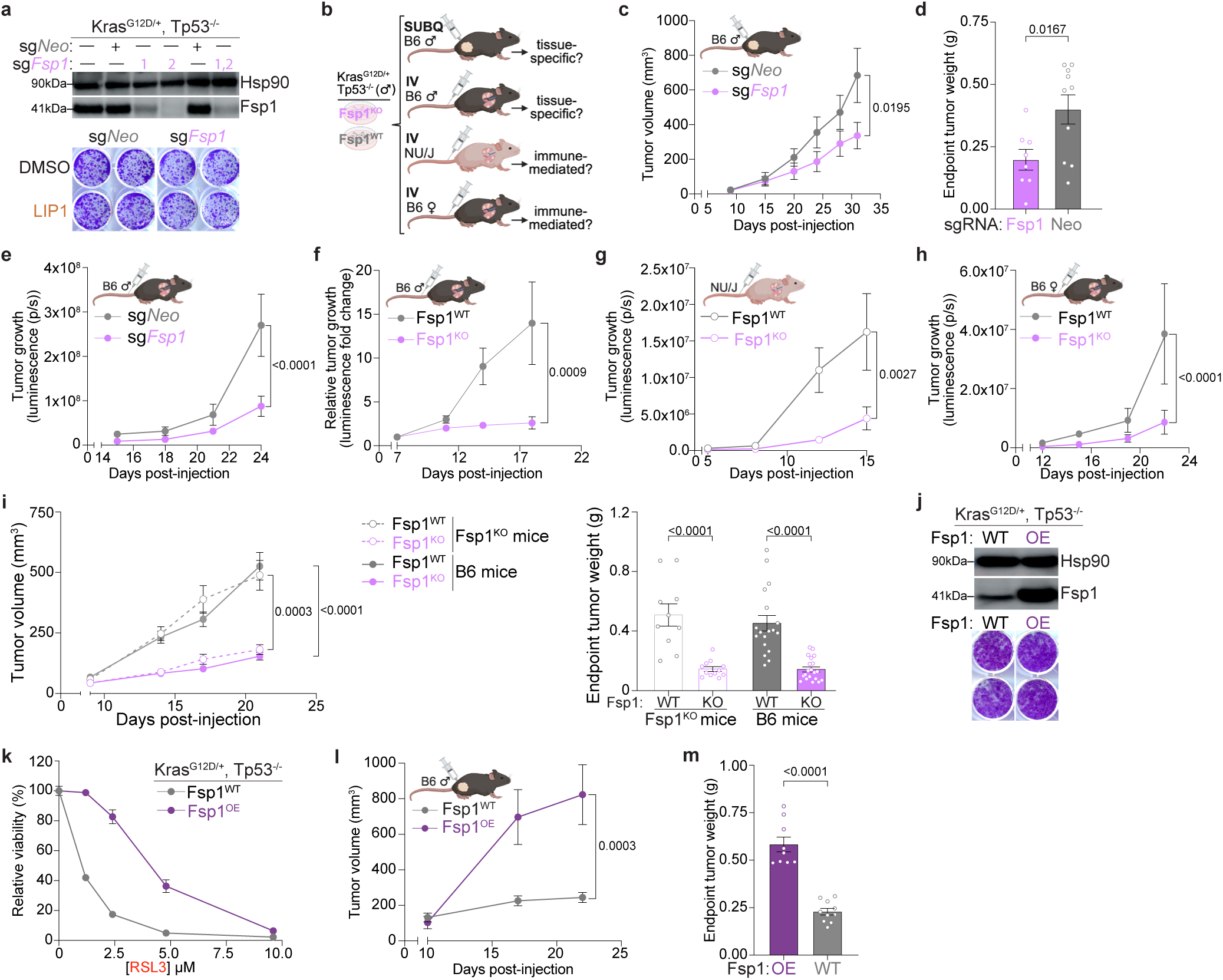
Fsp1 is required for cell-autonomous tumor growth *in vivo*. **(a)** Top: Western blot of KP LUAD cells with CRISPR/Cas9-mediated genetic deletion of Fsp1 with either two individual or duplexed sgRNAs. Bottom: Representative images of crystal violet clonogenic assay of KP LUAD cells with knockout of either control (Neo) or Fsp1. Cells were treated with 100nM LIP1. **(b)** Schematic depicting all transplantation models performed using isogenic KP, Fsp1^KO^ and Fsp1^WT^. **(c, d)** Longitudinal growth and endpoint tumor weights of KP sg*Fsp1* (n=8) versus control (sg*Neo* n=10) subcutaneous (subQ) xenograft tumor transplanted into C57BL/6J male mice. **(e)** Longitudinal lung tumor growth (measured via bioluminescence) in C57BL/6J male mice orthotopically transplanted with KP LUAD cells with CRISPR/Cas9-mediated knockout of Fsp1 (n=8) or control (n=7). **(f)** Longitudinal lung tumor growth (measured via bioluminescence normalized to first timepoint (day 7)) in C57BL/6J male mice orthotopically transplanted with isogenic KP, Fsp1^KO^ (n=6) and Fsp1^WT^ (n=7) cells. **(g)** Longitudinal lung tumor growth (measured via absolute bioluminescence) in NU/J immunocompromised mice orthotopically transplanted with isogenic KP, Fsp1^KO^ (n=6) and Fsp1^WT^ (n=7) cells. **(h)** Longitudinal lung tumor growth (measured via absolute bioluminescence) in C57BL/6J female mice with orthotopic transplantation of isogenic KP, Fsp1^KO^ (n=8) and Fsp1^WT^ (n=8) cells. **(i)** Longitudinal tumor growth and endpoint tumor weights of either isogenic KP, Fsp1^KO^ or Fsp1^WT^ subcutaneous (subQ) xenograft tumors transplanted in either C57BL/6J WT (Fsp1^KO^, male n=10 female n=10; Fsp1^WT^, male n=8, female n=10) or C57BL/6J *Fsp1*-knockout mice (Fsp1^KO^, males n=12, females n=12; Fsp1^WT^, male n=10, female n=10). **(j)** Western blot of KP LUAD cells with wildtype (WT) or overexpression (OE) of Fsp1. Representative images of crystal violet clonogenic assay of KP LUAD cells with Fsp1^WT^ or Fsp1^OE^. **(k)** CellTiter-Glo Luminescence viability assay of KP, Fsp1^WT^ or Fsp1^OE^ cells (5 biological replicates per group) with increasing concentrations of RSL3. **(l, m)** Longitudinal growth and endpoint weights of KP Fsp1^WT^ (n=10) or Fsp1^OE^ (n=10) subcutaneous (subQ) xenograft tumor transplanted into C57BL/6J male mice. Data are represented as mean values, error bars represent SEM, significance determined via one-way ANOVA with multiple comparisons (panel i), two-way ANOVA with Tukey’s multiple comparisons test (panels c, e, f, g, h, l), or two-sided student’s t-test (panels d, m). For gel source data, see Supplementary Data 1.

**Extended Data Fig. 5:**
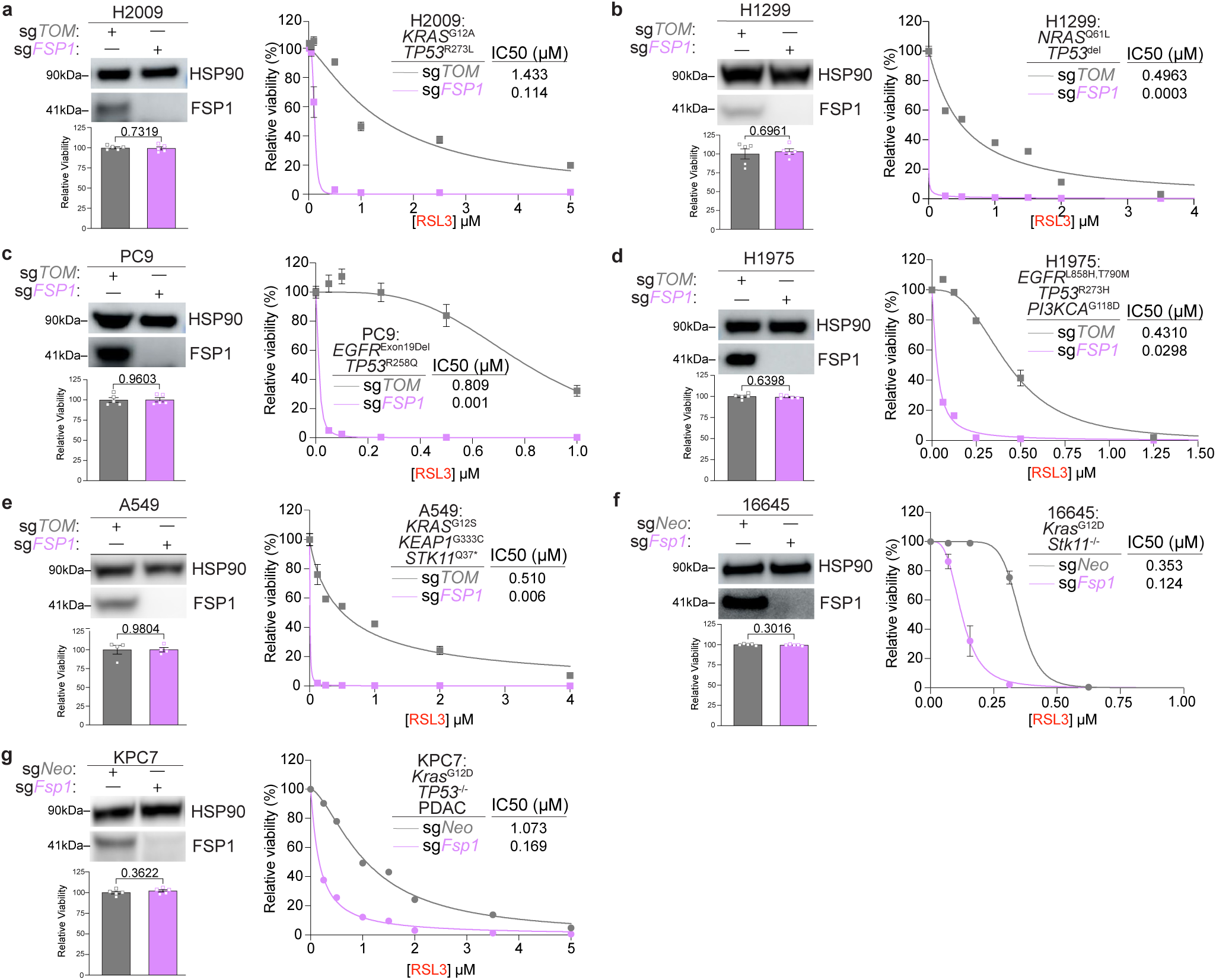
FSP1 loss *in vitro* does not induce ferroptosis without RSL3. **(a)** Top: Western blot; Bottom: baseline growth (normalized to control); and Right: CellTiter-Glo Luminescence viability assay of H2009 cells with the lentiviral addition of Cas9 and guide RNAs targeting FSP1 or a non-targeting control (TOM) (at least 4 biological replicates per group). Viability assay is upon increasing concentrations of RSL3. **(b)** As in (a), but with H1299 cells. **(c)** As in (a), but with PC9 cells. **(d)** As in (a), but with H1975 cells. **(e)** As in (a), but with A549 cells. **(f)** As in (a), but with 16645 cells. **(g)** As in (a), but with KPC7 cells. Data are represented as mean values, error bars represent SEM, significance determined via two-sided student’s t-test (panels a-g). For gel source data, see Supplementary Data 1.

**Extended Data Fig. 6:**
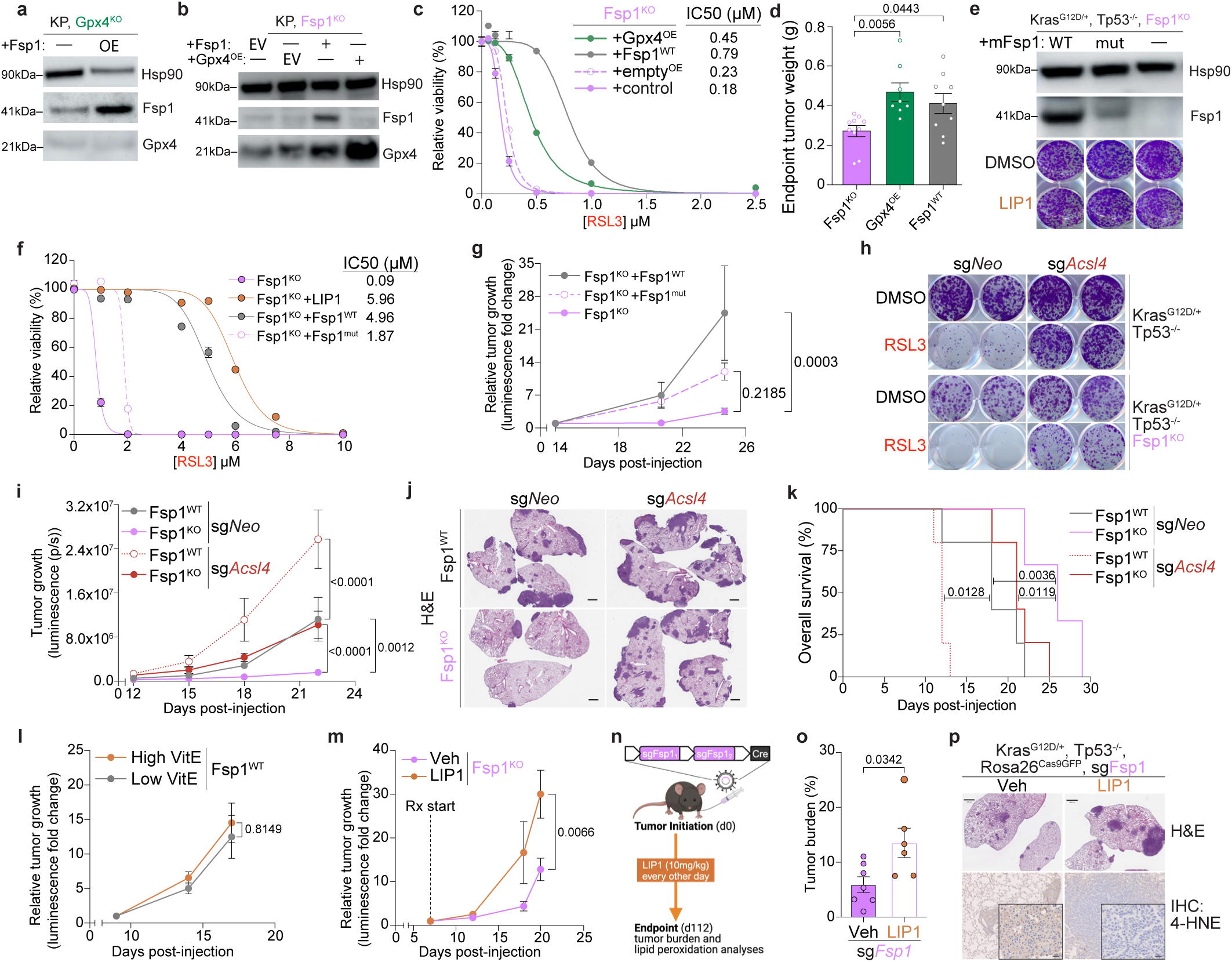
Fsp1 loss promotes tumor cell ferroptosis. **(a)** Western blot of KP, Gpx4^KO^ LUAD cells with either empty vector (—) or Fsp1^OE^. **(b)** Western blot of KP, Fsp1^KO^ LUAD cells with re-expression of either empty vector (—), Fsp1^WT^, Gpx4^OE^. **(c)** CellTiter-Glo Luminescence viability assay of KP, Fsp1^KO^ LUAD cells in (b) upon increasing concentrations of RSL3. **(d)** Endpoint tumor weights of subcutaneous (subQ) xenograft tumors from indicated cell lines transplanted in C57BL/6J female mice (control n=10; Fsp1^WT^, n=9; Gpx4^OE^; n=8). **(e)** Top: Western blot of KP, Fsp1^KO^ LUAD cells with re-expression of either empty vector (—), wildtype (WT) Fsp1, or mutant (mut) Fsp1. Bottom: Representative images of crystal violet clonogenic assay. **(f)** CellTiter-Glo Luminescence viability assay of KP, Fsp1^KO^ LUAD cells with re-expression of either empty vector (Fsp1^KO^), Fsp1^WT^, or Fsp1^mut^ upon increasing concentrations of RSL3. LIP1 used at 100nM. **(g)** Longitudinal tumor growth (measured via bioluminescence normalized to initial imaging timepoint (day14)) in C57BL/6J male mice orthotopically transplanted with isogenic KP, Fsp1^KO^ LUAD cells with re-expression of either empty control vector (Fsp1^KO^, n=7), Fsp1^WT^ (n=7), or Fsp1^mut^ (n=7). **(h)** Representative images of crystal violet clonogenic assay of KP, Fsp1^WT^ or Fsp1^KO^ LUAD cells with CRISPR/Cas9-mediated knockout of control (Neo) or Acsl4. **(i)** Longitudinal tumor growth (measured by bioluminescence) of C57BL/6J male mice with orthotopic transplantation of isogenic KP, Fsp1^KO^ and Fsp1^WT^ cells with CRISPR/Cas9-mediated knockout of control (Neo) or Acsl4 (WT, n=5; Acsl4^KO^, n=4; Fsp1^KO^, n=7; Fsp1^KO^Acsl4^KO^, n=7). **(j)** Representative H&E of experiment in (i). Scale bars: 1000µm. **(k)** Overall survival of C57BL/6J male mice with orthotopic transplantation of isogenic KP, Fsp1^KO^ and Fsp1^WT^ cells with CRISPR/Cas9-mediated knockout of control (Neo) or Acsl4 (n=5 per group). **(l)** Longitudinal tumor growth (measured via bioluminescence normalized to first imaging timepoint (day7)) of C57BL/6J mice orthotopically transplanted with KP, Fsp1^WT^ cells and receiving high (n=6) or low (n=5) VitE diets *ad libitum* 5 days pre-tumor initiation. **(m)** Tumor bioluminescence signal normalized to first imaging timepoint (day7) of C57BL/6J mice orthotopically transplanted with KP, Fsp1^KO^ cells and receiving Liproxstatin-1 (LIP1, n=7) or Vehicle (Veh, n=5) daily after tumor establishment. **(n)** Schematic of KP LUAD GEMMs intratracheally infected with pUSEC lentiviruses containing dual sgRNAs targeting Fsp1. Mice were dosed with LIP1 (n=6) or Vehicle (Veh, n=6) every other day starting from tumor initiation. **(o)** Tumor burden of KP LUAD GEMMs with CRISPR/Cas9-mediated knockout of Fsp1 and treated with Veh or LIP1. **(p)** Representative H&E and 4-HNE IHC of KP LUAD GEMM tumors with knockout of Fsp1 and treated with Veh or LIP1. Scale bars: 1000µm. Data are represented as mean values, error bars represent SEM, significance determined via one-way ANOVA (panel d), two-way ANOVA with Tukey’s multiple comparisons (panels g, i, l, m), two-sided student’s t-test (panel o), or Kaplain-Meier simple survival analysis (panel k). For gel source data, see Supplementary Data 1.

**Extended Data Fig. 7:**
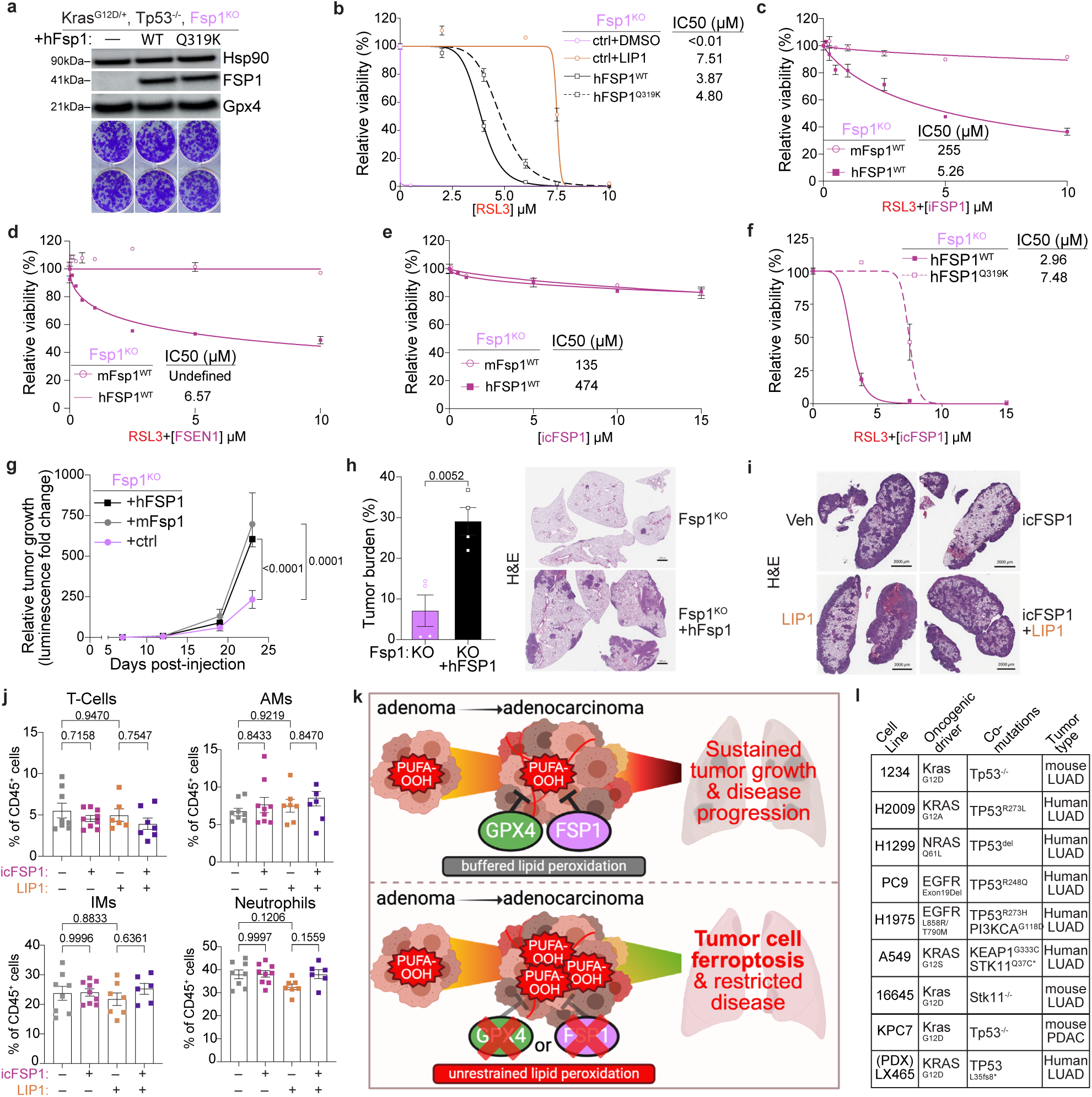
icFSP1 exerts on-target tumor-suppressive effect. **(a)** Top: Western blot of KP, Fsp1^KO^ LUAD cells with re-expression of either empty vector (—), wildtype (WT) hFSP1, or Q319K-mutant hFSP1. Bottom: Representative images of crystal violet clonogenic assay. **(b)** Cell-TiterGlo Luminescence viability assay of KP, Fsp1^KO^ LUAD cells with re-expression of either empty vector (—), WT hFSP1, or Q319K-mutant hFSP1. **(c)** CellTiter-Glo Luminescence viability assay of human or mouse Fsp1^WT^-expressing cells upon RSL3 (0.2µM) plus increasing concentrations of iFSP1. **(d)** CellTiter-Glo Luminescence viability assay as in (c) but with FSEN1. **(e)** CellTiter-Glo Luminescence viability assay of human or mouse Fsp1^WT^-expressing cells upon increasing concentrations of icFSP1. **(f)** CellTiter-Glo Luminescence viability assay of WT or Q319K-mutant hFSP1-expressing cells upon RSL3 (0.2µM) plus increasing concentrations of icFSP1. **(g)** Longitudinal tumor growth (measured via bioluminescence normalized to first imaging timepoint (day7)) in C57BL/6J mice orthotopically transplanted with isogenic KP, Fsp1^KO^ cells with re-expression of either human FSP1 (hFSP1, n=5), mouse Fsp1 (mFsp1, n=6), or control (n=6) vector. **(h)** Tumor burden and representative H&E of KP, Fsp1^KO^ (n=4) and hFSP1^WT^ (n=4) orthotopic lung tumors. Scale bars: 1000µm. **(i)** Representative H&E of KP, hFSP1^WT^ orthotopic lung tumors from indicated treatment groups. Scale bar: 2000µm. **(j)** Percent of T cells, alveolar macrophages, interstitial macrophages, and neutrophils of total CD45+ cells in tumor bearing lungs from mice treated with the indicated compounds for two weeks (Veh, n=7; LIP1, n=7; icFSP1, n=9; icFSP1+LIP1, n=6). **(k)** Graphical summary depicting ferroptosis as a barrier to lung cancer, as loss of either Fsp1 or Gpx4 induces tumor cell ferroptosis and restricts disease progression. **(l)** Summary table of cell lines, their respective tumor lineage, and mutations where Fsp1 loss resulted in tumor suppression *in vivo*. For all Cell-TiterGlo assays, n=5 biological replicates per group. Data are represented as mean values, error bars represent SEM, significance determined via one-way ANOVA with multiple comparisons (panels g, j) or student’s t test (panel h). For gel source data, see Supplementary Data 1.

## Methods References

51 Winslow, M.M., et al. Suppression of lung adenocarcinoma progression by Nkx2-1. Nature 473, 101–104 (2011).

52. Criscuolo, A., et al. Analytical and computational workflow for in-depth analysis of oxidized complex lipids in blood plasma. Nature Communications 13, 6547 (2022).

53. Goracci, L., et al. Lipostar, a Comprehensive Platform-Neutral Cheminformatics Tool for Lipidomics. Analytical Chemistry 89, 6257–6264 (2017).

54. Adams, K.J., et al. Skyline for Small Molecules: A Unifying Software Package for Quantitative Metabolomics. Journal of Proteome Research 19, 1447–1458 (2020).

55. Kang, Y.P., Kim, T.H., Ngoc Nguyen, C.T., Kim, S.M. & Kwon, S.W. Robust Determination of Coenzyme Q10 Redox Status Using Two Isotope-Labeled Internal Standards. Available at SSRN 4982487.

56. Pillai, R., et al. Glutamine antagonist DRP-104 suppresses tumor growth and enhances response to checkpoint blockade in KEAP1 mutant lung cancer. Sci Adv 10, eadm9859 (2024).

57. Created in BioRender. Vaughan, A. (2025) https://BioRender.com/99qhixq

## REFERENCES

1. Dixon, S.J., et al. Ferroptosis: an iron-dependent form of nonapoptotic cell death. Cell 149, 1060–1072 (2012).

2. Viswanathan, V.S., et al. Dependency of a therapy-resistant state of cancer cells on a lipid peroxidase pathway. Nature 547, 453–457 (2017).

3. Hangauer, M.J., et al. Drug-tolerant persister cancer cells are vulnerable to GPX4 inhibition. Nature 551, 247–250 (2017).

4. Berndt, C., et al. Ferroptosis in health and disease. Redox Biol 75, 103211 (2024).

5. Nakamura, T. & Conrad, M. Exploiting ferroptosis vulnerabilities in cancer. Nat Cell Biol (2024).

6. Friedmann Angeli, J.P., et al. Inactivation of the ferroptosis regulator Gpx4 triggers acute renal failure in mice. Nat Cell Biol 16, 1180–1191 (2014).

7. Yang, W.S., et al. Regulation of ferroptotic cancer cell death by GPX4. Cell 156, 317–331 (2014).

8. Bersuker, K., et al. The CoQ oxidoreductase FSP1 acts parallel to GPX4 to inhibit ferroptosis. Nature 575, 688–692 (2019).

9. Doll, S., et al. FSP1 is a glutathione-independent ferroptosis suppressor. Nature 575, 693–698 (2019).

10. Wiernicki, B., et al. Excessive phospholipid peroxidation distinguishes ferroptosis from other cell death modes including pyroptosis. Cell Death Dis 11, 922 (2020).

11. Conrad, M. & Pratt, D.A. The chemical basis of ferroptosis. Nat Chem Biol 15, 1137–1147 (2019).

12. Dixon, S.J. & Olzmann, J.A. The cell biology of ferroptosis. Nat Rev Mol Cell Biol 25, 424–442 (2024).

13. Kraft, V.A.N., et al. GTP Cyclohydrolase 1/Tetrahydrobiopterin Counteract Ferroptosis through Lipid Remodeling. ACS Cent Sci 6, 41–53 (2020).

14. Soula, M., et al. Metabolic determinants of cancer cell sensitivity to canonical ferroptosis inducers. Nat Chem Biol 16, 1351–1360 (2020).

15. Garcia-Bermudez, J., et al. Squalene accumulation in cholesterol auxotrophic lymphomas prevents oxidative cell death. Nature 567, 118–122 (2019).

16. Qiu, B., et al. Phospholipids with two polyunsaturated fatty acyl tails promote ferroptosis. Cell 187, 1177–1190 e1118 (2024).

17. Kagan, V.E., et al. Oxidized arachidonic and adrenic PEs navigate cells to ferroptosis. Nat Chem Biol 13, 81–90 (2017).

18. Doll, S., et al. ACSL4 dictates ferroptosis sensitivity by shaping cellular lipid composition. Nat Chem Biol 13, 91–98 (2017).

19. Dixon, S.J., et al. Human Haploid Cell Genetics Reveals Roles for Lipid Metabolism Genes in Nonapoptotic Cell Death. ACS Chem Biol 10, 1604–1609 (2015).

20. Mishima, E., et al. A non-canonical vitamin K cycle is a potent ferroptosis suppressor. Nature 608, 778–783 (2022).

21. Seiler, A., et al. Glutathione peroxidase 4 senses and translates oxidative stress into 12/15-lipoxygenase dependent- and AIF-mediated cell death. Cell Metab 8, 237–248 (2008).

22. Zilka, O., et al. On the Mechanism of Cytoprotection by Ferrostatin-1 and Liproxstatin-1 and the Role of Lipid Peroxidation in Ferroptotic Cell Death. ACS Cent Sci 3, 232–243 (2017).

23. Jiang, X., Stockwell, B.R. & Conrad, M. Ferroptosis: mechanisms, biology and role in disease. Nat Rev Mol Cell Biol 22, 266–282 (2021).

24. Nakamura, T., et al. Phase separation of FSP1 promotes ferroptosis. Nature 619, 371–377 (2023).

25. Eaton, J.K., et al. Selective covalent targeting of GPX4 using masked nitrile-oxide electrophiles. Nat Chem Biol 16, 497–506 (2020).

26. Yang, W.S. & Stockwell, B.R. Synthetic lethal screening identifies compounds activating iron-dependent, nonapoptotic cell death in oncogenic-RAS-harboring cancer cells. Chem Biol 15, 234–245 (2008).

27. Xavier da Silva, T.N., Schulte, C., Alves, A.N., Maric, H.M. & Friedmann Angeli, J.P. Molecular characterization of AIFM2/FSP1 inhibition by iFSP1-like molecules. Cell Death Dis 14, 281 (2023).

28. Hendricks, J.M., et al. Identification of structurally diverse FSP1 inhibitors that sensitize cancer cells to ferroptosis. Cell Chem Biol 30, 1090–1103 e1097 (2023).

29. Nakamura, T., et al. Integrated chemical and genetic screens unveil FSP1 mechanisms of ferroptosis regulation. Nat Struct Mol Biol 30, 1806–1815 (2023).

30. Yoshioka, H., et al. Identification of a Small Molecule That Enhances Ferroptosis via Inhibition of Ferroptosis Suppressor Protein 1 (FSP1). ACS Chem Biol 17, 483–491 (2022).

31. Sanchez-Rivera, F.J., et al. Rapid modelling of cooperating genetic events in cancer through somatic genome editing. Nature 516, 428–431 (2014).

32. Ding, H., et al. Activation of the NRF2 antioxidant program sensitizes tumors to G6PD inhibition. Sci Adv 7, eabk1023 (2021).

33. Lignitto, L., et al. Nrf2 Activation Promotes Lung Cancer Metastasis by Inhibiting the Degradation of Bach1. Cell 178, 316–329.e318 (2019).

34. Romero, R., et al. Keap1 loss promotes Kras-driven lung cancer and results in dependence on glutaminolysis. Nat Med 23, 1362–1368 (2017).

35. Best, S.A., et al. Distinct initiating events underpin the immune and metabolic heterogeneity of KRAS-mutant lung adenocarcinoma. Nat Commun 10, 4190 (2019).

36. Takahashi, N., et al. 3D Culture Models with CRISPR Screens Reveal Hyperactive NRF2 as a Prerequisite for Spheroid Formation via Regulation of Proliferation and Ferroptosis. Mol Cell 80, 828–844 e826 (2020).

37. Zou, Y., et al. A GPX4-dependent cancer cell state underlies the clear-cell morphology and confers sensitivity to ferroptosis. Nat Commun 10, 1617 (2019).

38. Vidigal, J.A. & Ventura, A. Rapid and efficient one-step generation of paired gRNA CRISPR-Cas9 libraries. Nat Commun 6, 8083 (2015).

39. Ingold, I., et al. Selenium Utilization by GPX4 Is Required to Prevent Hydroperoxide-Induced Ferroptosis. Cell 172, 409–422 e421 (2018).

40. Cancer Genome Atlas Research, N. Comprehensive molecular profiling of lung adenocarcinoma. Nature 511, 543–550 (2014).

41. Muller, F., et al. Elevated FSP1 protects KRAS-mutated cells from ferroptosis during tumor initiation. Cell Death Differ 30, 442–456 (2023).

42. Koppula, P., et al. A targetable CoQ-FSP1 axis drives ferroptosis- and radiation-resistance in KEAP1 inactive lung cancers. Nat Commun 13, 2206 (2022).

43. Zavitsanou, A.M., et al. KEAP1 mutation in lung adenocarcinoma promotes immune evasion and immunotherapy resistance. Cell Rep 42, 113295 (2023).

44. Wu, K., El Zowalaty, A.E., Sayin, V.I. & Papagiannakopoulos, T. The pleiotropic functions of reactive oxygen species in cancer. Nat Cancer 5, 384–399 (2024).

45. Cheung, E.C. & Vousden, K.H. The role of ROS in tumour development and progression. Nat Rev Cancer 22, 280–297 (2022).

46. Yant, L.J., et al. The selenoprotein GPX4 is essential for mouse development and protects from radiation and oxidative damage insults. Free Radic Biol Med 34, 496–502 (2003).

47. Mei, J., Webb, S., Zhang, B. & Shu, H.B. The p53-inducible apoptotic protein AMID is not required for normal development and tumor suppression. Oncogene 25, 849–856 (2006).

48. Nguyen, H.P., et al. Aifm2, a NADH Oxidase, Supports Robust Glycolysis and Is Required for Cold- and Diet-Induced Thermogenesis. Mol Cell 77, 600–617 e604 (2020).

49. Drijvers, J.M., et al. Pharmacologic Screening Identifies Metabolic Vulnerabilities of CD8(+) T Cells. Cancer Immunol Res 9, 184–199 (2021).

50. Matsushita, M., et al. T cell lipid peroxidation induces ferroptosis and prevents immunity to infection. J Exp Med 212, 555–568 (2015).

